# VC1 catalyzes a key step in the biosynthesis of vicine from GTP in faba bean

**DOI:** 10.1101/2020.02.26.966523

**Authors:** Emilie Björnsdotter, Marcin Nadzieja, Wei Chang, Leandro Escobar-Herrera, Davide Mancinotti, Deepti Angra, Hamid Khazaei, Christoph Crocoll, Albert Vandenberg, Frederick L. Stoddard, Donal M. O’Sullivan, Jens Stougaard, Alan H. Schulman, Stig U. Andersen, Fernando Geu-Flores

## Abstract

Faba bean is a widely adapted and high-yielding legume cultivated for its protein-rich seeds^1^. However, the seeds accumulate the anti-nutritional pyrimidine glucosides vicine and convicine, which can cause haemolytic anaemia—favism—in the 400 million individuals genetically predisposed by a deficiency in glucose-6-phosphate dehydrogenase^2^. Here, we identify the first enzyme associated with vicine and convicine biosynthesis, which we name VC1. We show that *VC1* co-locates with the major QTL for vicine and convicine content and that the expression of *VC1* correlates highly with vicine content across tissues. We also show that low-vicine varieties express a version of *VC1* carrying a small, frame-shift insertion, and that overexpression of wild-type *VC1* leads to an increase in vicine levels. *VC1* encodes a functional GTP cyclohydrolase II, an enzyme normally involved in riboflavin biosynthesis from the purine GTP. Through feeding studies, we demonstrate that GTP is a precursor of vicine both in faba bean and in the distantly related plant bitter gourd. Our results reveal an unexpected biosynthetic origin for vicine and convicine and pave the way for the development of faba bean cultivars that are free from these anti-nutrients, providing a safe and sustainable source of dietary protein.

## Main Text

According to the UN’s Intergovernmental Panel on Climate Change (IPCC), switching to a plant-based diet can reduce carbon emissions, especially in the West ^3^. The suggested change in diet will require a wider and more varied cultivation of locally adapted protein crops. On a worldwide basis, faba bean (**Fig. 1a**) has the highest yield of the legumes after soybean (1.92 Mg/ha in 2013-2017) ^4^ and the highest seed protein content of the starch-containing legumes (29% dry-matter basis) ^5^. Furthermore, faba bean is adapted to cool climates such as Mediterranean winters and northern European summers, where soybean performs poorly ^6^. The main factor restricting faba bean cultivation and consumption is the presence of the anti-nutritional compounds vicine and convicine (**Fig. 1b**). Already in the 5^th^ century BC, the Greek philosopher Pythagoras discouraged his followers from eating faba bean seeds, warning against a potentially fatal outcome ^7^. Indeed, faba bean ingestion may trigger favism**—**haemolytic anaemia from faba beans**—**in the 400 million individuals genetically predisposed to it (∼4% of the world population). These individuals display a deficiency in glucose-6-phosphate dehydrogenase, which is common in regions with historical endemic malaria and renders red blood cells susceptible to oxidative challenges. Vicine and convicine themselves are not strong oxidizing agents, but their metabolic products**—**divicine and isouramil**—**can cause irreversible oxidative damage in red blood cells leading to haemolysis (**Fig. 1b**). In contrast to the well-described aetiology of favism ^2^, the biosynthetic pathway of vicine and convicine in faba bean remains obscure.

**Figure 1.**
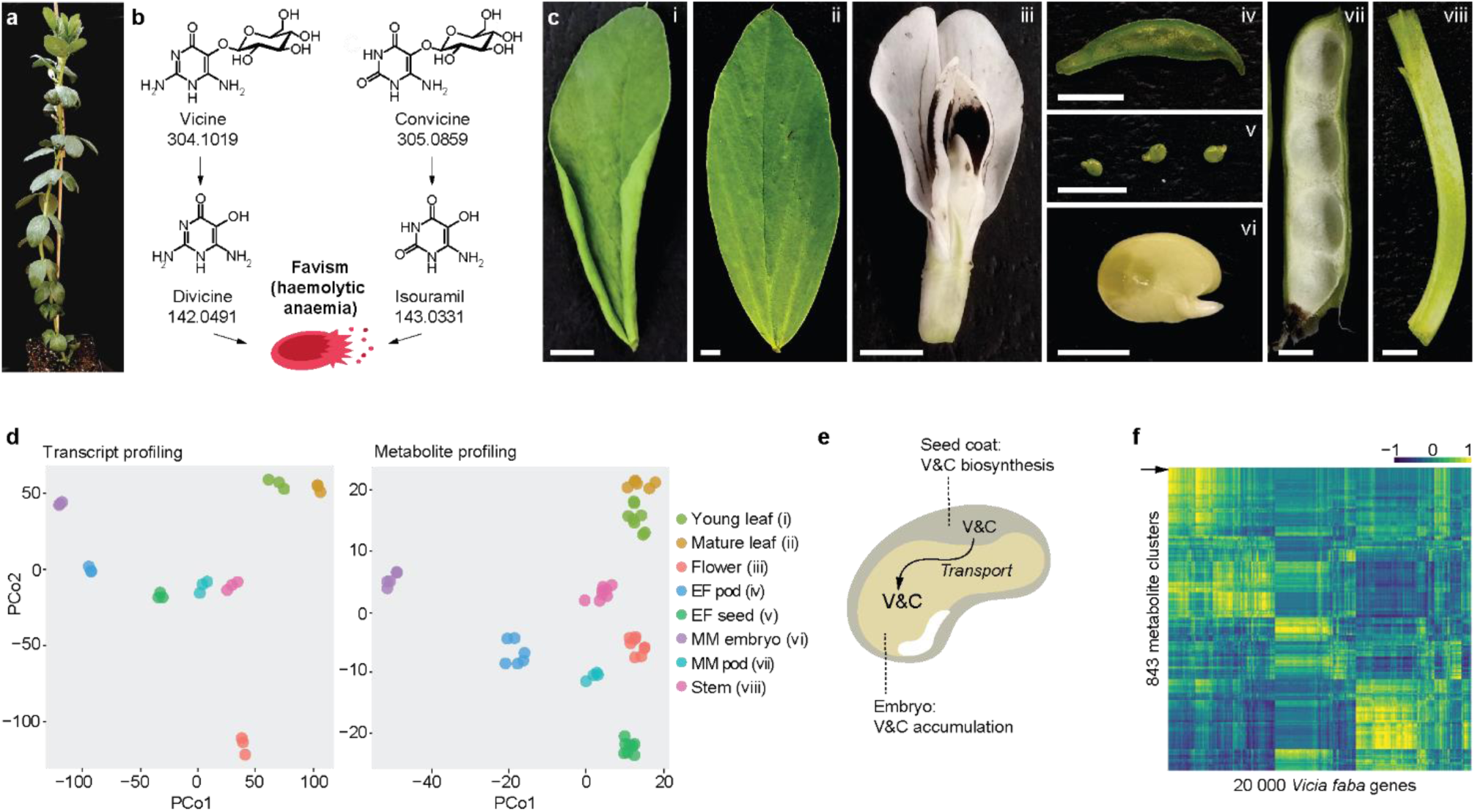
Gene expression analysis and metabolite profiling of eight faba bean tissues. **(a)** Faba bean plant at the onset of flowering. **(b)** The effect of vicine and convicine in individuals affected by favism. Once ingested, vicine and convicine are hydrolysed to divicine and isouramil, respectively. These metabolic products cause irreversible oxidative stress in red blood cells, leading to favism - haemolytic anaemia. Exact neutral masses are shown below compound names. **(c)** Faba bean tissues used for the gene expression and metabolite profiling: i) young leaves, ii) mature leaves, iii) flowers, iv) whole seeds at an early seed-filling stage (EF seeds), v) pods from an early seed-filling stage (EF pods), vi) embryos at mid maturation stage (MM embryo), vii) pods at the mid maturation stage (MM pods), viii) stems. Scale bars correspond to 5 mm. **(d)** Principal coordinate analysis of the gene expression and metabolite profiling datasets. Samples corresponding to the same tissue cluster together. All tissues are represented by distinct clusters. See tissue abbreviations above. **(e)** Current hypothesis on the translocation of vicine and convicine from biosynthetic, maternal tissues (e.g. seed coat) to the embryo. V&C, vicine and convicine. **(f)** Heat map representing the correlations of 843 metabolite clusters with 20 000 faba bean genes. MM embryos were not included in this analysis. The arrowhead indicates the metabolite feature cluster representing vicine (cluster 103).

In order to uncover genes associated with the biosynthesis of vicine and convicine in faba bean, we carried out a combined gene expression analysis and metabolite profiling of eight aerial tissues of the inbred line Hedin/2 (**Fig. 1c**). For the gene expression analysis, we assembled the raw RNA-seq data consisting of both short and long reads into a high-quality transcriptome composed of 49,277 coding sequences (**Extended Data Table 1**) (**Supplementary File 1**). We then mapped the short reads from each tissue onto the coding sequences, thus generating an expression matrix (**Supplementary File 2**). For the metabolite profiling, we analysed methanolic extracts using reverse-phase liquid chromatography coupled to high-resolution mass spectrometry, which yielded 1,479 unique metabolic features. We arranged these features into 852 clusters, each composed of one or more metabolic features with matching retention times and similar abundance patterns across tissues (**Supplementary File 3**). Cluster 103 was composed of two features whose *m/z* values corresponded to protonated vicine (feature 89_ID; theoretical *m/z*: 305.1097; experimental *m/z*: 305.1099) and its cognate aglucone (protonated vicine aglucone; feature 108_ID; theoretical *m/z*: 143.0569; experimental *m/z*: 143.0567). We confirmed that this cluster represented vicine by analysing a commercial standard and observing the same two features at a similar retention time. In both the gene expression and the metabolite datasets, all tissues could be clearly distinguished from one another using principal coordinate analysis (**Figure 1d)**.

We then proceeded to analyse gene-to-metabolite correlations. The content of vicine and convicine in seeds is maternally determined ^8^, which suggests that vicine and convicine are synthesized in maternal tissues and transported from there to developing embryos (**Fig. 1e**). To account for the possibility of translocation, we excluded isolated embryos from the analysis and computed the Pearson correlation coefficients across the seven remaining tissues (**Fig. 1f**). We then looked closely at the 20 genes most tightly correlated with vicine as represented by cluster_103 (**Supplementary File 4**). Among them, *evg_1250620* stood out by showing the highest expression level in whole seeds (seed coats plus embryos) at an early seed-filling stage (Fig. 2a) ^9, 10^ The gene encoded an isoform of 3,4-dihydroxy-2-butanone-4-phosphate synthase/GTP cyclohydrolase II, a bifunctional enzyme normally involved in riboflavin biosynthesis (**Extended Data Fig. 1**). A fragment of this gene, mis-annotated as *reticuline oxidase-like*, was previously identified among five other gene fragments based on gene expression comparison between normal- and low-vicine and convicine cultivars ^11^. More recently, Khazaei et al. (2017) ^12^ showed that a SNP coding for a silent mutation within this fragment distinguished between normal- and low-vicine and convicine cultivars in a diversity panel of 51 faba bean accessions.

**Figure 2.**
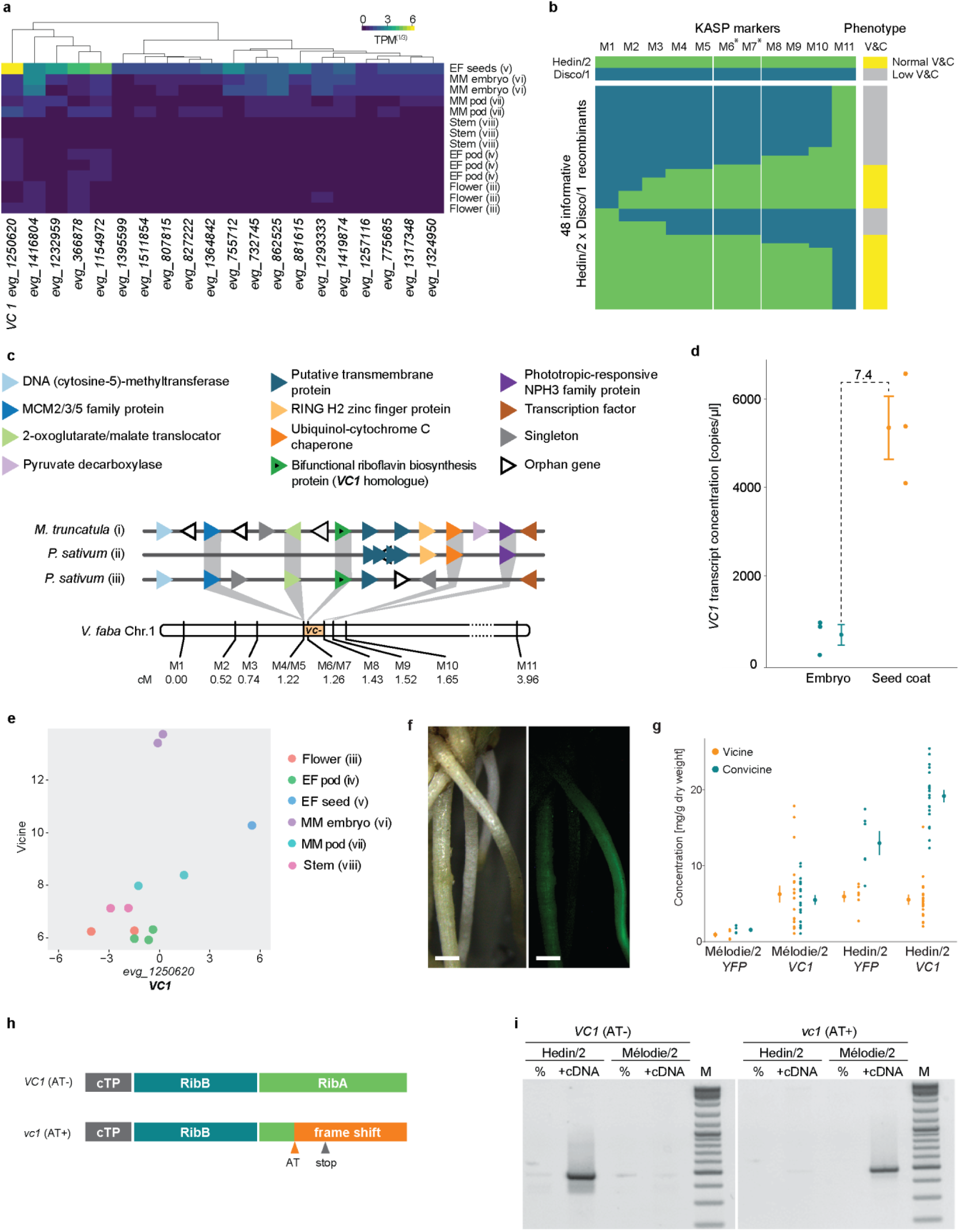
Identification of *VC1* as the *vc-* gene. **a)** Expression profile of the 20 genes most tightly correlated with vicine accumulation. The gene with the highest expression in whole seeds at an early maturation stage (EF seeds) is *evg_1250620* (*VC1*). None of these genes had detectable expression levels in leaf samples. **(b)** Narrowing of the genetic *vc-*interval using a Hedin/2 (normal-vicine and convicine) × Disco/1 (low-vicine and convicine) fine mapping population. Genotypes were assessed using competitive allele-specific PCR (KASP) markers. The genotypes and phenotypes of the parent lines are colour-coded and shown at the top. Allele calls and phenotypes of 48 informative recombinants are shown below using the same colour coding. Markers with an asterisk are positioned within the *VC1* gene. Marker sequences are described in **Extended Data Table 2**, and vicine and convicine levels are shown in **Extended Data Fig. 2**. V&C, vicine and convicine. **(c)** Syntenic context view of the alignment between the *V. faba vc*^-^ interval and collinear segments of *M. truncatula* (i - chr 2 from 1,801,324 to 1,875,086 bp) and *Pisum sativum* (ii - chr1 from 364,253,845 to 364,332,337 bp, iii - chr1 from 364,630,606 to 364,960,000 bp). Protein-coding genes are shown as differently coloured triangles, where triangles of the same colour represent a group of orthologous genes. Gene annotations are taken from the *M. truncatula* assembly Mt4.0v2. The genetic distance between markers on chromosome 1 of *V. faba* (Chr1) is shown in centimorgans (cM). **(d)** *VC1* transcript abundance in embryo and seed coat of the normal-vicine line Hedin/2 as determined by ddPCR. For each tissue, the individual data points represent biological variation, where each data point is the average of three technical replicates. Error bars represent the overall standard deviation per tissue. **(e)** Correlation between the logarithms of vicine content (metabolic feature 89) and *VC1* transcript abundance across Hedin/2 tissues as shown by the initial gene expression analysis and metabolite profiling. **(f)** Hairy roots of faba bean transformed with *YFP* under the control of the *pLjUbi* promoter. Pictures taken under white light (left) and UV light (right) are shown. The scale bar corresponds to 1 mm. **(g)** Vicine and convicine content in hairy roots transformed with *YFP* (control) or *VC1* under the control of the *pLjUbi* promoter in the background of either Mélodie/2 (low-vicine and convicine) and Hedin/2 (normal-vicine and convicine) lines. Error bars represent standard deviation. **(h)** Predicted functional domains of VC1 and the effect of the AT dinucleotide insertion (AT) in *vc1*. cTP, chloroplast transit peptide; RibB, 3,4-dihydroxy-2-butanone-4-phosphate synthase domain, RibA, GTP cyclohydrolase II domain. **(i)** Selective PCR amplification of *VC1* from Hedin/2 seed coat cDNA and *vc1* from Mélodie/2 seed coat cDNA. No cDNA was added to the negative controls (%). M, size marker.

All known low-vicine and convicine cultivars are derived from a single genetic source. The low vicine trait is inherited as a single recessive locus, termed *vc*^-^, but the causal gene remains unknown ^8^. Previous work had placed the *vc*^-^ locus within a 3.6 cM interval on chromosome 1 ^13^. We greatly refined the genetic interval carrying *vc*^-^ to 0.21 cM by mapping the low-vicine and convicine phenotype in a population of 1,157 pseudo F2 individuals from a cross between normal-(Hedin/2) and low-vicine and convicine (Disco/1) inbred lines (**Fig. 2b-c**). Within an overall context of conserved micro-colinearity, *vc*^-^ was bounded by markers defining an approximately 52-kb interval containing only eight genes in the genome of *Medicago truncatula* (*Medtr2g009220* to *Medtr2g009340*, corresponding to chr2:1,834,249-1,886,637). One of these eight *Medicago* genes, *Medtr2g009270*, encodes an isoform of 3,4-dihydroxy-2-butanone-4-phosphate synthase/GTP cyclohydrolase II (**Fig. 2c**). Moreover, the SNP identified by Khazaei et al. ^12^ and a second, independent SNP within *evg_1250620* co-segregated fully with the low-vicine and convicine phenotype, indicating that *evg_125620* is present within the refined 0.21-cM *vc*^-^ interval (**Fig. 2b**). Together with the gene-to-metabolite correlation results presented above, these genetic mapping results make *evg_1250620* a prime candidate for the *vc*^-^ gene. From here on, we will refer to *evg_1250620* as *VC1*.

In our gene expression profiling, *VC1* displayed high expression levels in whole seeds and low expression levels in isolated embryos (**Fig. 2a**). Because whole seeds are composed of seed coats and embryos, we hypothesized that *VC1* was highly expressed in seed coats, which are of maternal origin. In order to verify this, we conducted an additional gene expression study comparing seed coats to embryos of Hedin/2 using droplet digital PCR (ddPCR). This revealed that the expression of VC1 was around 7.4 times higher in seed coats than in embryos (**Fig. 2d**). It is worth noting that, in our combined gene expression and metabolite profiling, embryos stood out as having the highest vicine content, while showing only a moderate *VC1* gene expression (**Fig. 2e**). These results are consistent with the hypothesis that vicine and convicine are mainly synthesized in the seed coat and are transported to the embryo (**Fig. 1e**) ^9^ and suggest that *VC1* catalyses a key step in vicine biosynthesis.

We then investigated whether *VC1* was able to rescue the low-vicine and convicine phenotype. In the absence of an efficient transformation method for faba bean ^14^, we adopted a hairy root transformation protocol based on *Agrobacterium rhizogenes* ^*15*^. *We found that the ubiquitin* promoter *from Lotus japonicus* (*pLjUbi*) ^16^ could successfully drive the expression of *YFP* in hairy roots **(Fig. 2f)**, and that hairy roots of the normal-vicine line Hedin/2 accumulated several-fold more vicine and convicine than hairy roots of the low-vicine and convicine line Mélodie/2 (**Fig. 2g**). Transformation of Mélodie/2 hairy roots with the *VC1* coding sequence from Hedin/2 (also under the control of *pLjUbi*) led to a 7-fold increase in vicine levels compared to the *YFP* control, reaching the same levels as in the Hedin/2 *YFP* control. At the same time, a 3-fold increase in convicine levels was observed, reaching half the values of the Hedin/2 *YFP* control (**Fig. 2g**). Hairy roots of Hedin/2 transformed with *VC1* did not accumulate more vicine than the Hedin/2 *YFP* control, but the levels of convicine increased by a factor of 1.5 (**Fig. 2g**). The fact that *VC1* is able to complement the low-vicine and convicine phenotype of Mélodie/2 in hairy roots supports the hypothesis that *VC1* is the causal gene associated with the *vc*^-^ locus.

Next, we looked into the causal mutation leading to the low-vicine and convicine phenotype. First, we examined *VC1* expression in the seed coat, where *VC1* from Hedin/2 had shown high expression. Based on ddPCR, the expression level of *VC1* in Mélodie/2 was 4.7-times lower than in Hedin/2. This difference is not commensurate with the much lower vicine and convicine levels in Mélodie/2 (typically 10- to 40-times lower compared to Hedin/2 seeds). We then examined the *VC1* coding sequences cloned from seed coat cDNA. The coding sequence from Hedin/2 matched the sequence derived from our RNA-seq data exactly. In contrast, the sequence from Mélodie/2, which we designate *vc1*, contained a 2-nucleotide AT insertion causing a reading frame shift in the region encoding the GTP cyclohydrolase II (**Fig. 2h, Extended Data Fig. 2, Supplementary File 6**). Using seed coat cDNA and PCR primers able to distinguish between *VC1* and *vc1*, we detected only *VC1* in Hedin/2 whereas *vc1* was predominant in Mélodie/2 (**Fig. 2i)**. The AT insertion is located within the first half of the region encoding the GTP cyclohydrolase II and prevents the correct synthesis of at least half of the enzyme, including key residues that are necessary for activity ^17^ (**Fig. 2h, Extended Data Fig. 2**). This suggests that this AT insertion is the direct cause of the low vicine and convicine levels of Mélodie/2 (and all other known low-vicine and convicine cultivars) and that the GTP cyclohydrolase II domain of *VC1* is involved in the biosynthesis of vicine and convicine.

Vicine and convicine are pyrimidine glucosides and were thought to be derived from the orotic acid pathway of pyrimidine biosynthesis (**Fig. 3a**) ^18^. This is not consistent with our identification of *VC1*, which is presumably involved in purine-based riboflavin biosynthesis. Of the two putative enzymes encoded by the bifunctional *VC1*, GTP cyclohydrolase II catalyzes the first step of the riboflavin pathway, which is the conversion of the purine nucleoside triphosphate GTP into the unstable intermediate 2,5-diamino-6-ribosylamino-4(3*H*)-pyrimidinone 5’-phosphate (DARPP). Next, a deaminase converts DARPP into a second unstable intermediate, 5-amino-6-ribosylamino-2,3(1H,3H)-pyrimidinedione 5’-phosphate (ARPDP). We noticed a structural similarity between DARPP/ARPDP and vicine/convicine, respectively. Accordingly, we hypothesize that vicine and convicine are derived respectively from DARPP and ARPDP via a parallel, 3-step biochemical transformation (**Fig. 3a**). The first of these proposed transformations is a hydrolysis that has recently been shown to be catalyzed by COG3236 in bacteria and plants ^19^. Only two more steps would be necessary to produce vicine and convicine: a deamination and a glucosylation (**Fig. 3a**).

**Figure 3.**
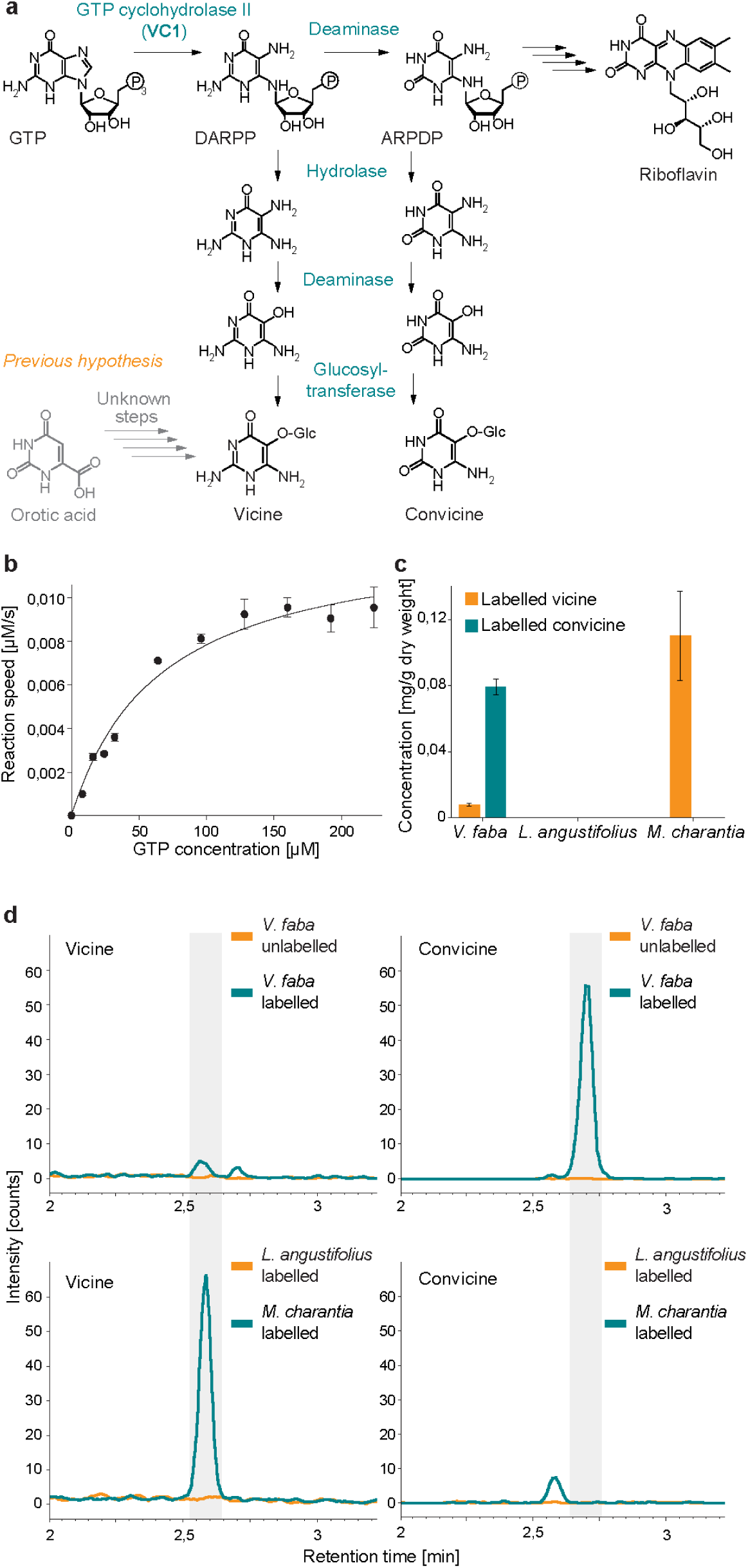
Characterization of VC1 as a GTP cyclohydrolase II involved in vicine and convicine biosynthesis and establishment of GTP as a biosynthetic precursor. **(a)** Proposed pathway for the biosynthesis of vicine and convicine. **(b)** Michaelis–Menten kinetics of the GTP to DARPP conversion catalyzed *in vitro* by purified VC1. **(c)** Feeding of *V. faba, L. angustifolius* and *M. charantia* roots with ^13^C_10_,^15^N_5_-GTP (labelled GTP) and its incorporation into vicine and convicine. Feeding with unlabelled GTP was performed as a control. **(d)** Elution profiles of labelled vicine (panels on the left) and labelled convicine (panels on the right) from the feeding experiments. The top row includes faba bean fed with labelled and unlabelled GTP. The bottom row includes *Lupinus angustifolius* (Fabaceae, non-vicine and convicine producer) and *Momordica charantia* (non-Fabaceae, vicine producer) fed with labelled GTP.

To test our pathway hypothesis, we first tested the activity of the VC1 protein *in vitro*. For this, we expressed a tagged version of VC1 in *E. coli* and purified it using affinity chromatography (**Extended Data Fig. 4a**). The purified enzyme was able to convert GTP to DARPP (**Extended data Fig. 4b**). Kinetic studies revealed a *K*_M_ value of 66 ± 12 µM and a turnover number of 1.6 ± 0.11 min^-1^ (**Fig. 3b**). These kinetic parameters resemble those of other functional GTP cyclohydrolase II enzymes ^20, 21, 22^. Then, we fed ^13^C_10_,^15^N_5_-GTP to Hedin/2 roots to determine whether GTP was a precursor for vicine and convicine. This resulted in the detection of both ^13^C_4_,^15^N_4_-vicine and ^13^C_4_,^15^N_3_-convicine, whereas the feeding of unlabelled GTP did not (**Fig. 3c-d**). We performed analogous feeding studies with narrow-leafed lupin (*Lupinus angustifolius*), a legume that does not accumulate vicine and convicine, and these did not result in the detection of labelled vicine and convicine (**Fig. 3c-d**). Finally, we fed ^13^C_10_,^15^N_5_-GTP to roots of bitter melon (*Momordica charantia*), which is a phylogenetically remote species (*Cucurbitaceae*) that accumulates vicine but not convicine. This resulted in the detection of the same labelled vicine species seen previously in faba bean (^13^C_4_,^15^N_4_-vicine) (**Fig. 3c-d**). These feeding experiments establish GTP as a precursor for vicine and convicine and indicate that vicine biosynthesis from GTP evolved independently at least twice.

In summary, we have identified *VC1* as a key gene in the biosynthesis of vicine and convicine as well as the mutated *vc1* gene that represents the single known genetic source of low vicine and convicine content. Our study also demonstrates that the pyrimidine glucosides vicine and convicine are not derived from pyrimidine metabolism but from purine metabolism, specifically from intermediates in the riboflavin pathway. This work represents a stepping stone towards the complete elucidation of the biosynthetic pathway of vicine and convicine as well as the full elimination of these anti-nutritional compounds from faba bean.

## Methods

### Gene expression analysis, metabolite profiling, and gene-to-metabolite correlations

#### Plant growth and sampling

Faba bean plants of the inbred line Hedin/2 were grown in the field at Sejet International ApS (Horsens, Denmark). The following tissue types were collected: i) young leaf (closest to the shoot meristem, not fully open); ii) mature leaf (fully open); iii) flower (banner petals open); iv) pod at early seed-filling (EF) stage; v) whole seed at EM stage (containing seed coat and embryo); vi) embryo at mid maturation (MM) stage; vii) pod at MM stage; viii) stem (4 - 5 cm segments positioned 5 cm below the top of the shoot meristem). Sample collection was carried out at the same time of the day to reduce the influence of circadian rhythm. Tissue samples were harvested, flash frozen on site, and later ground and split into pools for RNA isolation and metabolite extraction. For EF seeds, due to prolonged dissection time resulting in small volume of samples difficult to split, six separate replicates were harvested, of which three were used for transcriptome analysis and another three for metabolite profiling. The ground tissue pools were stored at −80 °C until further analysis.

#### Gene expression analysis

Total RNA was extracted from ground tissues using the NucleoSpin RNA Plant extraction kit (Macherey-Nagel). Non-strand-specific cDNA libraries of 250-300 bp were synthesized and sequenced by Novogene (Hong Kong) using the HiSeq PE150 sequencer (Illumina), resulting in 30-43 million reads per sample. Additionally, two strand-specific Illumina libraries (Novogene, Hong Kong) and one PacBio library (Earlham Institute, UK) prepared from a pool of the RNA samples were sequenced, yielding 64 and 0.5 million reads, respectively. A *de novo* assembly of the *V. faba* Hedin/2 gene set was created using Trinity 2.4.0 ^1^. First, an assembly was made independently for each tissue. Triplicates were used alongside long reads from the PacBio dataset. For the pool of RNA samples, only two duplicates were employed. To reduce the redundancy within each assembly, the assemblies were subjected to CD-HIT-EST clustering with a sequence identity threshold of 0.95 and a word size of eight ^2^. The clustered assemblies were then merged into one to create a combined gene set. Next, the EvidentialGene pipeline was run using standard settings to filter for quality and further decrease for redundancy ^3^. The quality of the assemblies were accessed by mapping reads back to the assemblies using BWA-mem and BUSCO ^4, 5^. Transcript quantification was performed by using Bowtie2, R and RSEM ^6, 7^. Bowtie2 was run in the following modes: no discordant, no gaps in the first 1000 bases, no-mixed, and end-to-end mode. Finally, the set of transcripts was filtered with an expression cut-off set to 1 transcript per million mapped reads (TPM) across the tissues.

#### Metabolite profiling

Ground tissues were freeze-dried and around 2.5 mg of dry material was extracted with 200 µl of 60% MeOH containing 50 µM caffeine as internal standard. The mixture was shaken for 15 min at 1 200 rpm and centrifuged at 13 500 × *g* for 5 min. The supernatant was diluted 10x with 15% MeOH and cleared through 0.22 µm filters. Reversed-phase LC-MS analysis was performed on a Thermo Fisher Dionex UltiMate 3000 RS HPLC/UHPLC system fitted with a Kinetex EVO C18 column (100 × 2.1 mm, 1.7 µm, 100 Å, Phenomenex) and interfaced to an ESI compact QqTOF mass spectrometer (Bruker). The eluent flow rate was 0.3 ml/min and the column temperature was kept constant at 40 °C. Mobile phases A and B consisted of 0.05% formic acid in water and 0.05% formic acid in acetonitrile, respectively. The elution profile was 0 – 5 min, 0% B constant; 5 – 24 min, 0 – 100% B linear; 24 – 26 min, 100% B linear, 26 – 27 min, 100% – 0% B linear; 27 – 35 min, 100% B constant. ESI mass spectra were acquired in positive ionization mode with the following parameters: capillary voltage of 4500 V; end plate offset of −500 V; source temperature of 250 °C; desolvation gas flow of 8.0 l/min; nebulizer pressure of 2.5 bar. Nitrogen was used as desolvation and nebulizer gas. The scanned *m/z* range was 50 to 1000. Sodium formate clusters were used for internal mass calibration and were introduced at the beginning of each run (first 0.5 min). Each tissue extract was injected twice (technical replicates) and a blank sample was run every 10 injections. The raw LC-MS chromatograms were mass calibrated, converted to mzXML format and submitted to XCMS Online (ver. 3.7.1) for alignment, feature detection and quantification ^8^. A multijob analysis was performed using the default settings for UPLC/Bruker Q-TOF instruments and considering the following sample groups: EF pods (n = 8), MM pods (n = 4), EF seeds (n = 6), MM embryos (n = 6), flowers (n = 4), stems (n = 4), young leaves (n = 4), mature leaves (n = 3), and blanks (n = 8). Biological and technical replicates were treated as independent samples. Metabolite features were defined as mass spectral peaks of width between 5 and 20 seconds and signal-to-noise ratio of at least 6:1. Metabolic features derived from the mass calibrant (retention time < 0.5 min) were removed. The dataset was further filtered by removing metabolic features whose intensity in any of the tissue sample groups was not significantly different from that in the blank sample group (p < 0.01 in Student’s T-test). After filtering, the intensities of the remaining metabolite features were normalized to the dry weight of the samples and to the signal of the internal standard (the protonated molecular ion of caffeine). The normalized intensity profile of each metabolite feature was centred and scaled. Using MultiExperiment Viewer (ver. 4.9) ^9^, the metabolite features were subjected to complete-linkage hierarchical clustering analysis (HCA) based on the Pearson’s correlation coefficient between their centred and scaled intensity profiles. The HCA dendrogram was manually divided into discrete metabolic clusters of the largest possible height and composed entirely of metabolite features with overlapping median retention times (difference of < 6 s). As indicated in the main text, we identified a cluster (cluster 108) composed of two features, corresponding to protonated vicine (median *m/z* 305.1099) and protonated vicine aglucone (median *m/z* 143.0567). The separate running of a commercial vicine standard confirmed that these two features represented vicine. Two analogous metabolic features were found for convicine (median *m/z* 306.0994 and 144.0491). However, due to vicine and convicine having the same retention time in our experimental setup, these features represented not only the convicine-related [M+1] ions, but also the respective vicine-related [M+2] ions. Accordingly, these additional features were not investigated further.

#### Gene-to-metabolite correlations

Prior to calculating correlation coefficients, expression and metabolite data was normalized using Poisson-seq ^10^. We then used the ‘cor’ function of R (version 3.4.3) ^11^ to calculate the Pearson correlation coefficients for gene expression (quantified as TPM) versus the normalized intensity of metabolic features. The correlations obtained were then averaged across the metabolic features in each metabolic cluster. For all tissues except EF seeds, individual samples were directly matched in the correlation analysis. For EF seeds, separate samples were used for gene expression and metabolite profiling, and the mean of the replicates was used for the correlation analysis. Since vicine and convicine are likely to be produced in maternal tissues and transported to the embryo (see main text), MM embryos were excluded from the analysis. A total of 17 samples from the following tissues were used in this analysis: flowers (3), stems (3), young leaves (3), mature leaves (2), EF pods (3), EF seeds (1), and MM pods (2). See **Supplementary File 5** for full details and the R scripts used.

### ddPCR-based quantification of VC1 expression in embryo vs seed coat

Plants were grown in the greenhouse of the Viikki Plant Science Centre (Helsinki, Finland). Embryo and seed coat tissues were harvested from Hedin/2 plants at the mid maturation stage and flash frozen. Frozen tissues were ground using TissueLyser MM300 oscillatory mixer mill (Qiagen Retsch). For embryo tissue, RNA was extracted from single embryos using 1 ml TRIzol (Thermo Fisher Scientific) following the manufacturer’s instructions. The extracted RNA was treated with DNaseI (Ambion) and purified with an RNeasy MinElute Cleanup Kit (Qiagen). For seed coats, RNA was extracted from 100 mg of powdered tissue using the RNeasy Plant Mini Kit (Qiagen) including DNAse treatment. Extractions were made as three technical replicates per plant and as three plants for each tissue. First-strand cDNA was synthesized using Superscript IV reverse transcriptase (Invitrogen) and primed with oligo(dT). Droplet digital PCR was carried out on a QX200 AutoDG Droplet Digital PCR System (Bio-Rad). The PCR reaction contained 10 μL of 2x QX200 ddPCR EvaGreen Supermix (Bio-Rad), 100 nM forward primer (CTTCTTGCATTCTCCTCATTTCCTC) and 100 nM reverse primer (CCCTCCAGATACCAATGCAGCTTTAACC), 1 μl cDNA, and nuclease-free water to a final volume of 20 μL. The PCR program consisted of 95 °C for 5 min; 40 cycles of denaturation at 95 °C for 30 s followed by annealing/extension at 58 °C for 1 min (ramp rate of 2 °C s^-1^); and signal stabilization at 4 °C for 5 min. The resulting data were analyzed with QuantaSoft software v1.7 (Bio-Rad).

### Specific amplification of VC1 and vc1 from seed coat of Hedin/2 and Mélodie/2

Seed coat RNA was extracted and converted to cDNA as described in the previous section. For the specific amplification of *vc1* (with AT insertion), we used forward primer GACATATTTGGATCTGCCACATATG and reverse primer TCCTCAAAGACCAGTAGCACC. PCR was carried out using 1 μl cDNA using the following temperature program: 94 °C for 2 min; 40 cycles of denaturation at 94 °C for 30 sec, annealing at 58 °C for 30 sec, and extension at 72 °C for 40 sec; signal stabilization at 72 °C for 5 min. For the specific amplification of the active *VC1* form (no AT insertion), an alternative forward primer was used: GACATATTTGGATCTGCCACTTG. A similar amplification program was used, but with an annealing temperature of 54 °C.

### Targeted analysis of vicine and convicine

Approximately 2.5 mg of dry tissue was weighed, ground and extracted with 200 µl of 60% MeOH containing 8 µM uridine as internal standard. The mixture was shaken for 15 min at 1 200 rpm at room temperature, followed by a 5-min centrifugation at 12 000 rpm. The supernatant was diluted 10x with 90% acetonitrile and cleared using a 0.22-µm filter. HILIC chromatography coupled to mass spectrometry was used to detect vicine and convicine through a method developed by Purves *et al*. (Purves, 2018). Chromatography was performed on an Advance UHPLC system (Bruker, Bremen, Germany) with an Acquity UPLC BEH Amide column (2.1 × 50 mm, 1.7 µm, Waters). The mobile phases consisted of solvent A (10 mM ammonium acetate and 0.1% formic acid in water) and solvent B (10 mM ammonium acetate and 0.1% formic acid in 90:10 acetonitrile:water). The following gradient program was run at a flow rate of 400 µl/min: from 100% - 90% B for 0.5 min; from 90% to 75% B for 3.5 min; from 75% to 100% B for 0.2 min; 100% B for 3.8 min. The HILIC column was coupled to an EVOQ Elite triple quadrupole mass spectrometer (Bruker, Bremen, Germany) equipped with an electrospray ionisation source (ESI). The ion spray voltage was maintained at −5000 V. Cone temperature was set to 350 °C and cone gas pressure to 20 psi. The temperature of the heated probe was set to 275 °C and the probe gas pressure to 30 psi. Nebulizing gas was set to 40 psi and collision gas to 1.6 mTorr. Nitrogen was used as cone gas, probe gas and nebulizing gas and argon as collision gas. Multiple reaction monitoring (MRM) was carried out in negative mode and the transitions used were 303 → 141 for vicine (collision energy (CE) = 15 eV), 304 → 141 for convicine (CE = 19 eV), and 243 → 200 for uridine (CE = 6 eV). Uridine signals were used for normalization, and external standard curves (1-2000 nM) were used for quantification of vicine and convicine. Bruker MS Workstation software (Version 8.2.1, Bruker, Bremen, Germany) was used for data acquisition and processing.

### Fine mapping of vc^-^

Inbred lines Hedin/2 and Disco/1 (normal- and low-vicine phenotype, respectively) were crossed to obtain an F_2_ population of 73 F_2_ individuals. Selfed seeds from 39 F_3_ individuals, which were heterozygous across the previously defined *vc*^-^ interval ^12^, were grown to form a pseudo-F_2_ population of 1,157 individual plants segregating for the *vc*^-^ gene. Individual SNP (Single Nucleotide Polymorphism) KASP assays were selected from previous maps based on the 3.4-cM interval reported by Khazaei ^12^ or designed based on markers mined from RNA-seq data. The markers used are described in **Extended Data Table 2**. KASP markers developed by Webb et al. ^13^ bounding the *vc*^-^ interval described by Khazaei et al. ^12^ were initially used to screen the Hedin/2 x Disco/1 pseudo-F_2_ population for putative recombinants. 90 recombinants were found, which were then genotyped for the full panel of *vc*^-^-targeted polymorphisms together with the parental stocks. A genetic map fragment was constructed using R/QTL ^14^. Dry seeds of 48 informative recombinants were harvested, ground to flour, and analysed for vicine and convicine using the targeted analysis described above.

### *Cloning of VC1 and vc1* coding sequences

The *VC1* coding sequence was cloned from Hedin/2 roots as well as from seed coats. When using roots as starting material, we used 2-week-old seedlings grown on vermiculite at room temperature. We used the Spectrum Plant Total RNA Kit (Sigma-Aldrich) to extract RNA. cDNA was synthesized from RNA using the SuperScript™ III First-Strand Synthesis System (Thermo Fisher Scientific) and oligo (dT)_20_ primers. The coding sequence was amplified by PCR using cDNA as template and the following primers: ATGGCAGCTGCTACTTTCAAT and TCAAACAGTGATTTTAACACCATTGTTA. The PCR product was cloned into vector pJET1.2/blunt using CloneJet PCR Cloning Kit (Thermo Scientific) and sequenced. When using seed coats as starting material, RNA was extracted as described above for ddPCR and cloned as described below for *vc1*.

The *vc1* coding sequence was cloned from Melódie/2 seed coats harvested from greenhouse-grown plants. The seed coats were isolated 20-25 days after tripping (hand pollination). RNA was extracted from frozen seed coat powder as described above for ddPCR. First-strand cDNA was carried out also as described above for ddPCR. The coding sequence of *vc1* was amplified by PCR using cDNA as template as well as primers CTTCTTGCATTCTCCTCATTTCCTC (forward) and TCCTCAAAGACCAGTAGCACC (reverse), which target the 5’ and 3’ ends of the transcript, respectively. The PCR product was cloned into pGEM®-T (Promega) and sequenced.

### Overexpression of VC1 in hairy roots

In order to introduce 3 silent mutations that removed *BpiI* and *BsaI* restriction sites, we synthesized the coding sequence of *VC1* cloned from root cDNA (GeneScript). The synthesized sequence was PCR amplified using primers ATGAAGACGGAATGATGGCAGCTGCTACTTTCAAT and ATGAAGACGGAAGCTCAAACAGTGATTTTAACACC, which added GoldenGate overhangs for creating an SC module ^15^. The level-0 plasmid SC-*VC1* was created in a 20 μl reaction containing 100 ng of the gel-purified PCR product, 100 ng of the target pICH vector, 5 U of T4 ligase (Thermo Scientific), 2.5 U of BpiI (Thermo Scientific), and 2 μl of 10x T4 ligase buffer. The following temperature programme was used: 25x (37 °C for 3 min, 16 °C for 4 min), 65 °C for 5 min, and 80 °C for 5 min. The overexpression construct *LjUbi:VC1* (**Supplementary File 7**) was created in a 20 μl reaction containing 100 ng of each of the following plasmids: PU-LjUbi, SC-*VC1*, T-35s, and pIV10, as well as 5U of T4 ligase, 2.5 U of BsaI (New England BioLabs), and 2 μl 10x T4 ligase buffer.

Seeds of Mélodie/2 and Hedin/2 were surface-sterilized for 10 min on 0.5% sodium hypochlorite and subsequently rinsed 5 times with sterile water. The sterilized seeds were germinated on petri dishes lined with moist filter paper and transferred to magenta boxes containing moist vermiculite. Plants were grown at 21 °C with a photoperiod of 16/8 h. In parallel, plasmids *LjUbi:YFP* ^16^ *and LjUbi:VC1* were conjugated into *Agrobacterium rhizogenes* GV3101 using triparental mating ^17^. We then infected the *in-vitro-*grown plants with the transformed *A. rhizogenes* using a protocol adapted from Kereszt et al. ^18^. Briefly, seedlings that had produced two true leaves were wounded at the hypocotyls and inoculated with a high-density suspension of *A. rhizogenes*. Inoculated plants were incubated in the dark for 48 h and then grown at 21 °C with a photoperiod of 16/8 h for 3-4 weeks. Hairy root tissue was flash frozen in liquid nitrogen and freeze-dried for targeted vicine and convicine analysis.

### Stably labelled precursor feeding experiments

Seeds of faba bean (Hedin/2), narrow-leafed lupin (cv. Oskar, purchased from HR Smolice, Poland) and bitter gourd (purchased from Bjarne’s Frø og Planter, Denmark) were germinated on moist paper. 3-4-day seedlings were transferred to 2-ml Eppendorf tubes, where they were fed for 72 h with 1.5 ml of 1 mM ^13^C_10_,^15^N_5_-GTP in 5 mM Tris buffer at pH 7.2 through the roots. As controls, seedlings were fed with unlabelled GTP instead. The entire roots were cut from the seedlings, frozen in liquid nitrogen, and freeze-dried. The targeted analysis of labelled vicine and convicine was carried out as described above for unlabelled vicine and convicine, except for the MRM transitions used, which were 311 → 149 (CE = 15 eV) for labelled vicine (^13^C_4_,^15^N_4_-vicine) and 311 → 148 (CE = 19 eV) for labelled convicine (^13^C_4_,^15^N_3_-convicine). For quantification, labelled vicine and convicine were assumed to have the same ionization efficiencies as their unlabelled forms.

### Expression and purification of His-tagged VC1

We predicted the chloroplast transit peptide (cTP) of VC1 using TargetP online (version 2.0) ^19^. An *E. coli* codon-optimized version of *VC1* coding for an N-terminal His-tag and lacking the predicted cTP-coding region (**Supplementary File 7**) was synthesized (GenScript) and cloned into expression vector pET22b(+) using restriction sites NdeI and HindIII. The plasmid was transformed into ArcticExpress (DE3) RIL *E. coli* competent cells (Agilent Technologies) and protein expression was performed mainly as described by Hiltunen *et al*. ^20^. Cells were grown at 37 °C and 220 rpm in 750 ml of selective LB media up to an OD_600_ of 0.5-0.7. The culture was cooled on ice and subsequently induced by adding IPTG to a final concentration of 1 mM. Protein expression took place for 24 h at 13°C and 170 rpm. After pelleting, cells were resuspended in 2 ml of lysis buffer (50 mM Tris, 300 mM NaCl, 0.01% β-mercaptoethanol, 2 mM imidazole, pH 7.5), and 400 µl of 25x cOmplete EDTA-free protease inhibitor was added before adding 0.2 mg of lysozyme. Following a 1 h incubation on ice and subsequent sonication, the lysate was cleared by centrifugation at 17 000 × *g* and 4 °C for 25 min. The His-tagged protein was immediately purified from the cleared lysate using affinity chromatography with stepwise elution. The lysate was gently shaken with 0.5 ml of Ni-NTA agarose suspension (Qiagen) for 1 h at 4 °C and transferred to a filter column where the liquid was drained. The matrix was washed 3 times with 1.5 ml washing buffer (50 mM Tris, 300 mM NaCl, 0.01% β- mercaptoethanol, 5 mM imidazole, pH 7.5). Elution was carried out using 1 ml of four different elution buffers with different imidazole concentrations (50 mM Tris, 300 mM NaCl, and either 20 mM, 50 mM, 100 mM, or 250 mM imidazole, pH 7.5). The different eluate fractions were analysed by SDS-PAGE, which revealed that most of the heterologously expressed protein eluted in the fraction with 250 mM imidazole (**Extended Data Fig. 4)**. To remove imidazole and concentrate the protein, the 250 mM imidazole fraction was buffer-exchanged into storage buffer (20 mM Tris, 200 mM NaCl, 5% (v/v) glycerol, pH 8.0) using a 30K Amicon filter. The purified enzyme was assayed immediately or stored at −20 °C, which preserved enzyme activity. A typical yield of purified VC1 from a 750 ml culture was 6 mg. Protein concentration was estimated using the Pierce™ BCA Protein Assay Kit (ThermoFisher).

### Enzyme assays and kinetics

Enzyme activity was analysed as previously reported ^21, 22^. The reaction was carried out in 200 µl and contained 50 mM Tris at pH 8.0, 100 mM NaCl, 10 mM MgCl_2_ and GTP at concentrations varying from 0-244 µM. The reaction was started by adding 5 µg of purified VC1. Conversion of GTP to the product 2,5-diamino-6-β-ribosyl-4(3*H*)-pyrimidinone-5’-phosphate (DARPP) was monitored by measuring absorbance at 310 nm for 5 min using a microplate reader (Spectramax M5, Molecular Devices). The reaction rate was calculated using the extinction coefficient for DARPP (7.43 cm^-1^ mM^-1^) as previously reported ^21, 22^. The kinetic parameters *K*_M_ and *V*_max_ were calculated by non-linear regression to fit the data to the Michaelis-Menten equation using Sigmaplot v13.0.

## Acknowledgements

This work was supported by Innovation Fund Denmark grant number 5158-00004B; Academy of Finland decisions 298314 and 314961; UK Biotechnology and Biological Science Research Council award BB/P023509/1; VILLUM Foundation Project 15476; and Danish National Research Foundation grant DNRF99. We acknowledge the technical assistance of Anne-Mari Narvanto and Laura Vottonen^3,7^ as well as the bioinformatic support and analyses by Jaakko Tanskanen^8^.

## Author Contributions

FGF, SUA, and AHS conceived research plan; EB, MN, WC, LEH, DM, and DA carried out experiments and data analysis; HK, CC, DOS, and FLS provided instrumentation and resources; JS, DOS, AHS, AV, SUA, FLS, and FGF developed project design and acquired funding; JS coordinated the project; MN and SUA prepared figures; SUA and FGF wrote the manuscript with input from all authors.

**Supplementary information** is available for this paper.

## Extended data

**Extended Data Table 1.** Parameters of the *Vicia faba* Hedin/2 transcript assembly. Open reading frames (ORFs) were predicted using Transdecoder.

| Feature | Stat |
| --- | --- |
| Transcripts | 49277 |
| ORFs | 35663 |
| Total bases | 41144820 |
| GC content | 42.54% |
| N50 | 1314 |
| Median contig length | 501 |
| Average contig length | 835 |
| Complete BUSCO | 94.60% |
| Single BUSCO | 89.90% |
| Duplicated BUSCO | 4.70% |
| Missing BUSCO | 3.90% |
| Fragmented BUSCO | 1.50% |

**Extended Data Table 2.**
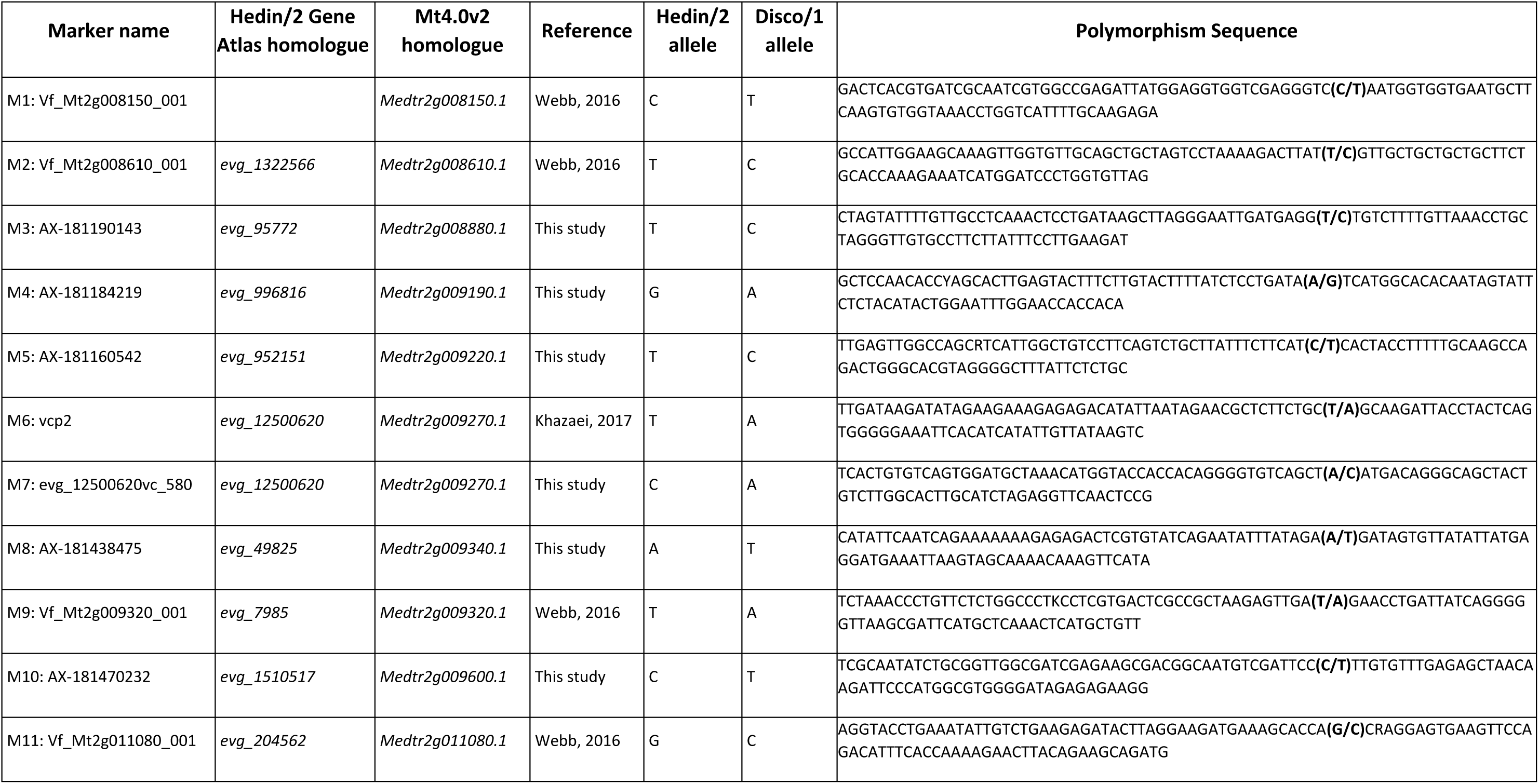
SNP KASP markers developed to saturate the *vc*^-^ interval. *V. faba* Hedin/2 and *M. truncatula* transcript IDs are listed. Full polymorphism sequences distinguishing between Hedin/2 and Disco/1 inbred lines are specified.

**Extended Data Figure 1.**
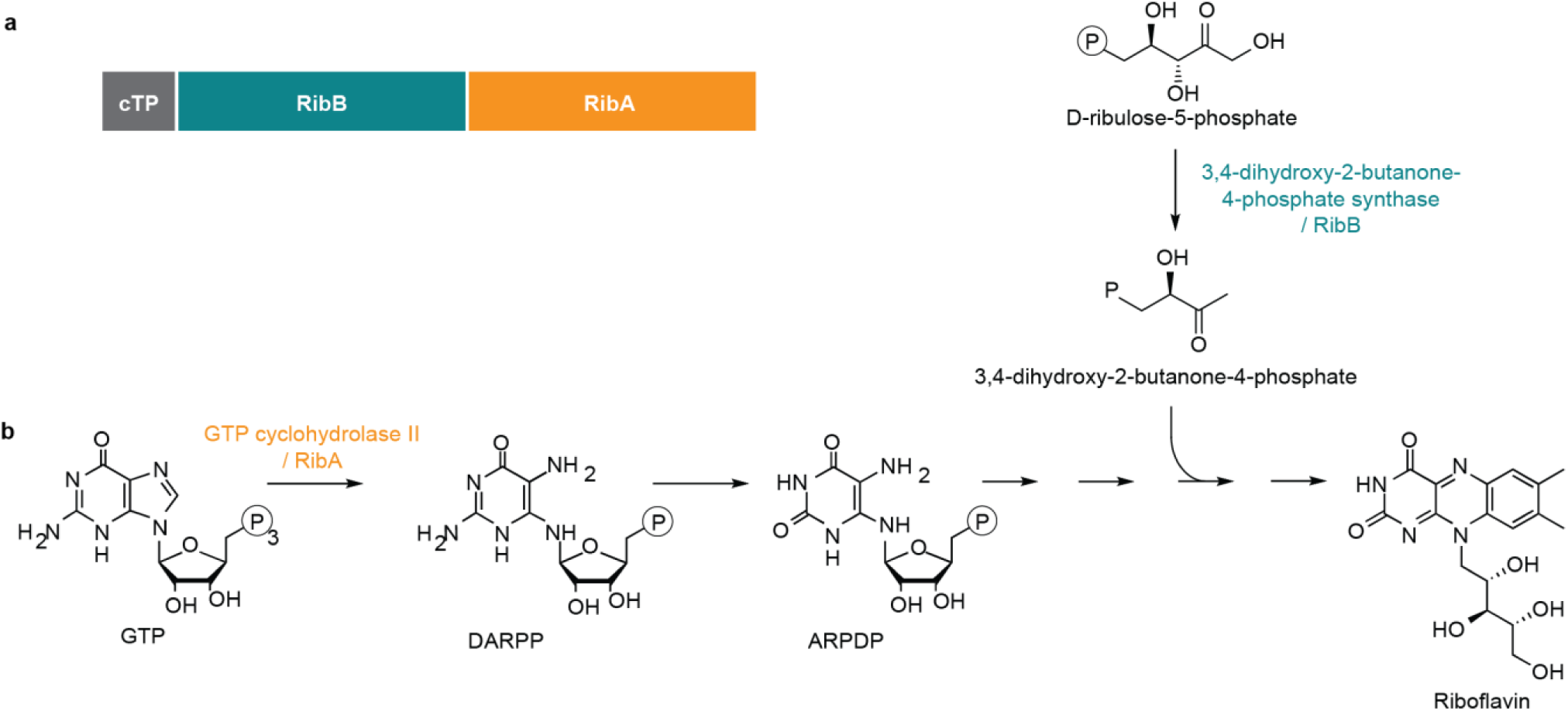
Canonical function of the bifunctional enzyme 3,4-dihydroxy-2-butanone 4-phosphate synthase/GTP cyclohydrolase II in plants. **(a)** Domain structure. The protein is composed of a chloroplast targeting peptide (cTP) fused to two catalytic domains: the 3,4-dihydroxy-2-butanone-4-phosphate synthase domain, also called RibB, and the GTP cyclohydrolase II domain, also called RibA. **(b)** Biochemical function of each catalytic domain in the context of the riboflavin biosynthesis pathway.

**Extended Data Figure 2.**
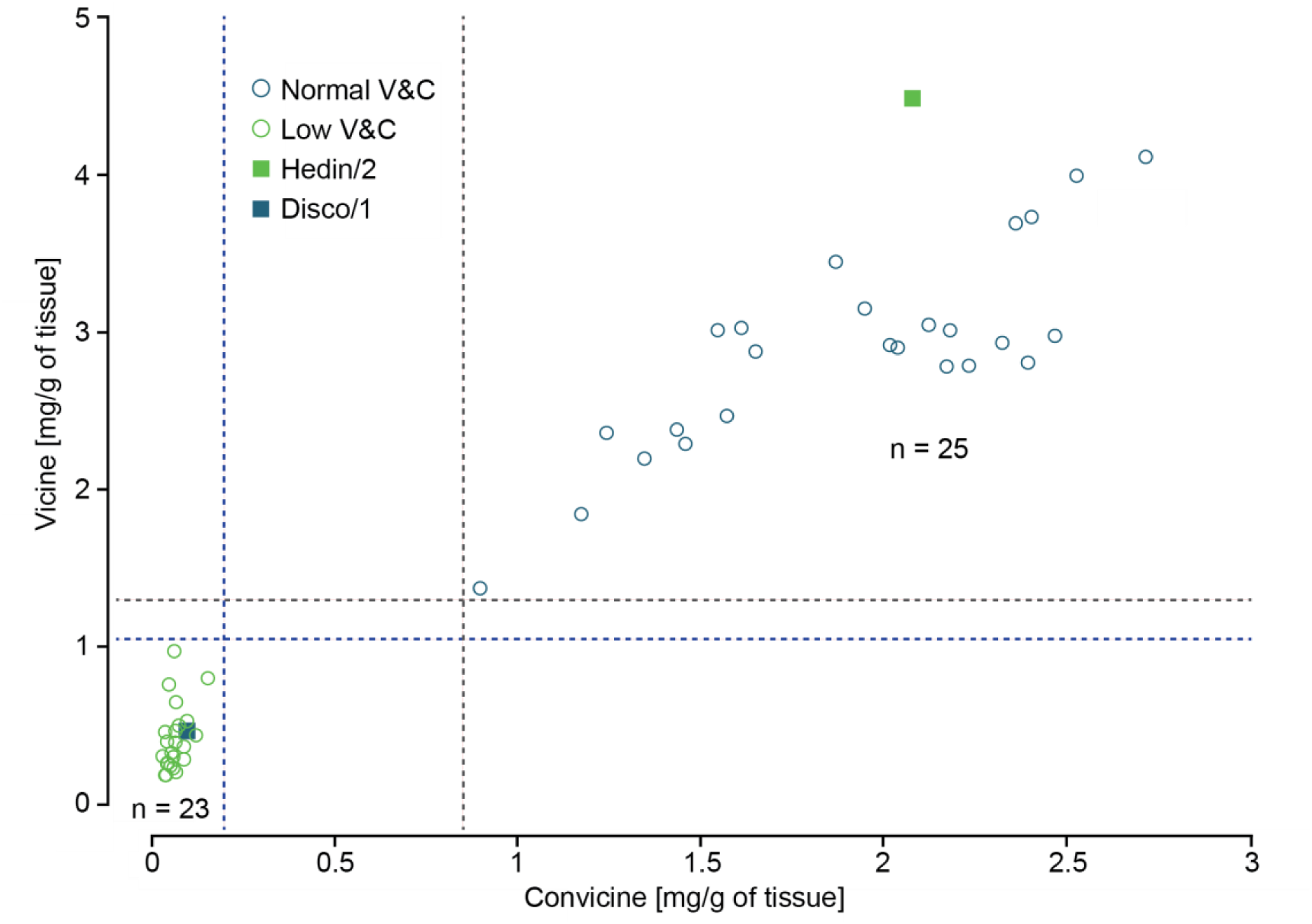
Seed vicine and convicine phenotypes of Hedin/2 × Disco/1 pseudo-F2 recombinants within the *vc*^-^ interval. Recombinants are classified as Normal (blue open circles) where vicine levels are >1.3 mg/g and convicine levels are >0.85 mg/g or as Low (green open circles) where vicine levels are <1.05 mg/g and convicine levels are <0.2 mg/g. Parental means are shown as squares for Hedin/2 (green) and Disco/1 (blue).

**Extended Data Figure 3.**
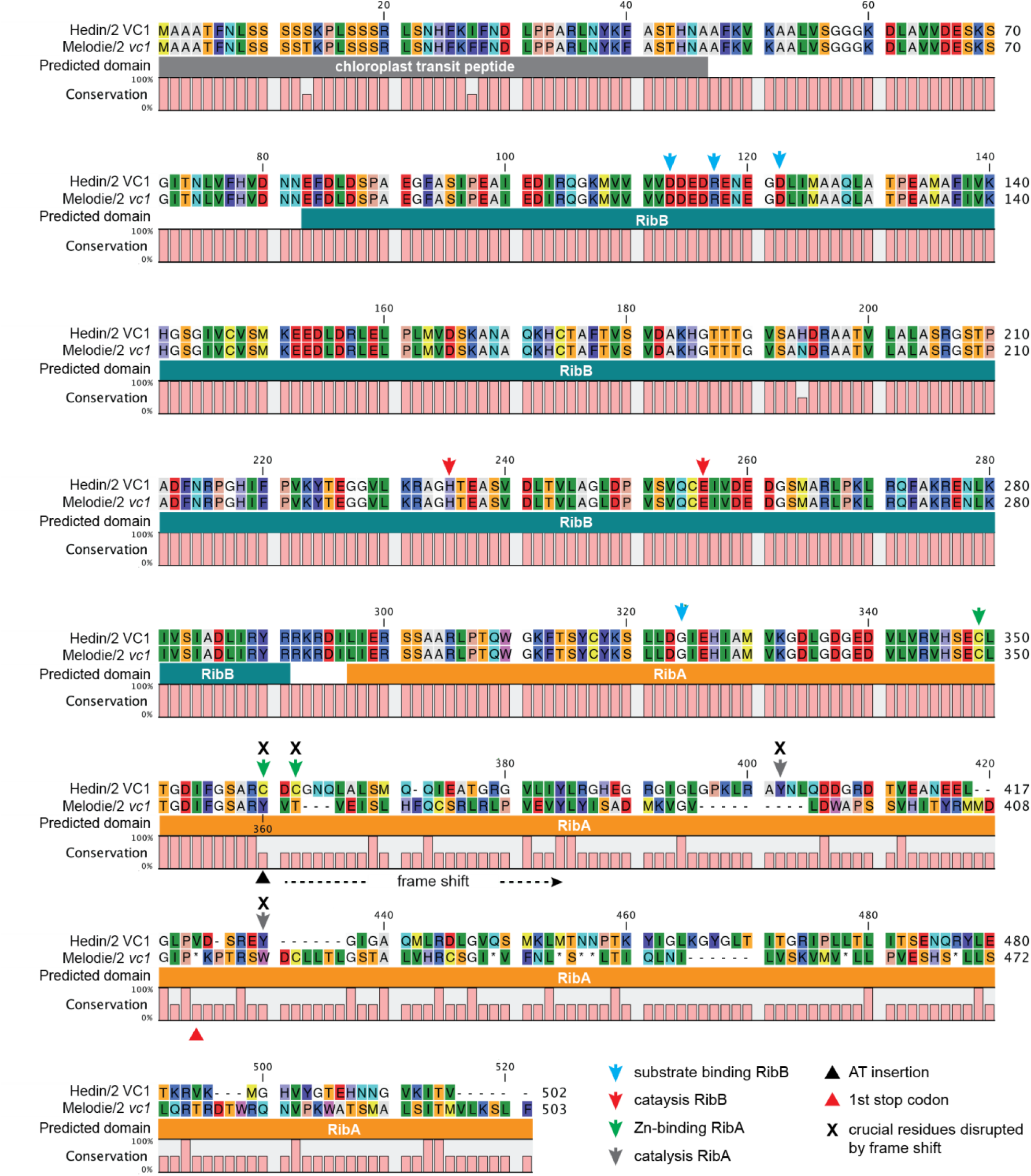
Consequence of the additional AT dinucleotide on the predicted VC1 protein. An alignment of Hedin/2 VC1 and Mélodie/2 vc1 amino acid sequences is shown. Predicted domains are shown underneath the alignment (RibB, 3,4-dihydroxy-2-butanone-4-phosphate synthase domain; RibA, GTP cyclohydrolase II domain). A measurement of residue conservation is shown underneath the predicted domains, distinguishing between identical residues and other scenarios (non-identical ones as well as gaps/insertions). The position of the AT insertion resulting in frame shift marked with a black triangle underneath the conservation score (position 360). The following key residues in VC1 enzymatic domains are marked: blue arrows, substrate binding in RibB; red arrows, catalysis in RibB; green arrows, Zn-binding cytosines in RibA; grey arrows, catalytic tyrosines in RibA. The residue prediction is based on Hiltunen et al. (2012).

**Extended Data Figure 4.**
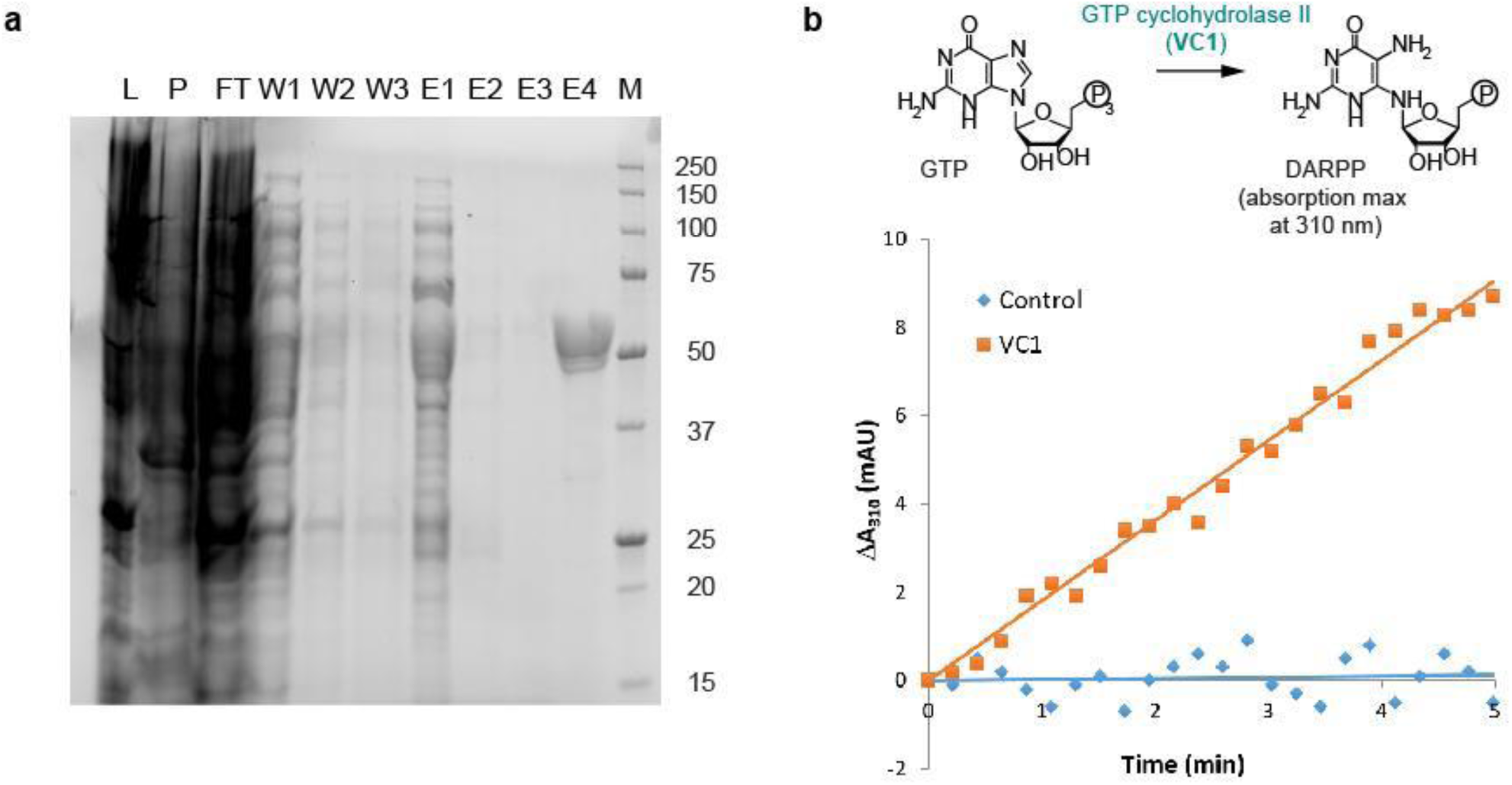
VC1 expression, purification, and assays. **(a)** SDS-PAGE gel showing the affinity-purification of His-tagged VC1 on a Ni-NTA matrix. L, lysate; P, pellet; FT, flow-through; W1-3, three consecutive wash fractions; E1-4, elutions with increasing concentration of imidazole (20, 50, 100, and 250 mM, respectively); M, molecular weight marker (given in kDa). The expected molecular weight of His-tagged VC1 was 51.3 kDa. After buffer exchange to remove the imidazole, fraction E4 was used for the subsequent assays. **(b)** Representative result of the GTP cyclohydrolase II assays measuring the appearance of DARPP, which presents an absorption maximum at 310 nm. The graph shows the increase in absorbance at 310 nm (ΔA_310_) against time for a control (no enzyme) and for an assay with purified VC1.

## SI Guide

**Supplementary File 1**. Transcriptome coding sequences in fasta format.

**Supplementary File 2**. Gene expression counts in transcripts per million (TPM).

**Supplementary File 3**. List of metabolic features, their grouping into clusters, and their abundances across tissue samples.

**Supplementary File 4**. List of top-20 genes correlated with vicine accumulation levels in all tissues except mid-maturation embryos.

**Supplementary File 5**. R scripts used to analyse gene-to-metabolite correlations. **Supplementary File 6**. *VC1* and *vc1* cDNA sequences and predicted amino acid sequences. **Supplementary File 7**. Design of the expression constructs used in the study.

## Main References

1. Duc, G. et al. Faba Bean. Grain Legumes 141–178 (2015) doi: 10.1007/978-1-4939-2797-5_5.

2. Luzzatto, L. & Arese, P. Favism and glucose-6-phosphate dehydrogenase deficiency. N Engl. J. Med. vol. 378 1068–1069 (2018).

3. Shukla, P.R. et al. The Intergovernmental Panel on Climate Change (IPCC). Summary for Policymakers. In: Climate Change and Land: an IPCC special report on climate change, desertification, land degradation, sustainable land management, food security, and greenhouse gas fluxes in terrestrial ecosystems. In press (2019).

4. FAOSTAT. Food and Agriculture Organization of the United Nations. http://www.fao.org/faostat/en/#home. Visited 27.02.2020.

5. Feedipedia - Animal Feed Resources Information System - INRA, CIRAD, AFZ and FAO. https://www.feedipedia.org. Visited 27.02.2020.

6. Stoddard, F.L. Grain legumes: an overview. Chapter 5, pp. 70–87 in: Legumes in Cropping Systems, eds. Murphy-Bokern, D., Stoddard, F.L., & Watson, C.A. CAB International, Oxford, UK (2017).

7. Meletis, J. & Konstantopoulos, K. Favism-from the ‘avoid fava beans’ of Pythagoras to the present. Haema 7, 17–21 (2004).

8. Duc, G., Sixdenier, G., Lila, M. & Furstoss, V. Search of genetic variability for vicine and convicine content in *Vicia faba* L.: a first report of a gene which codes for nearly zero-vicine and zero-convicine contents. in 1. International Workshop on Antinutritional Factors (ANF) in Legume Seeds, Wageningen (Netherlands), 23-25 Nov 1988(Pudoc, 1989).

9. Ramsay, G. & Griffiths, D. W. Accumulation of vicine and convicine in *Vicia faba* and *V. narbonensis*. Phytochemistry 42, 63–67 (1996).

10. Lin, J.Y. et al. Similarity between soybean and *Arabidopsis* seed methylomes and loss of non-CG methylation does not affect seed development. Proc. Natl. Acad. Sci. U. S. A. 114, E9730–E9739 (2017).

11. Ray, H., Bock, C. & Georges, F. Faba Bean: Transcriptome analysis from etiolated seedling and developing seed coat of key cultivars for synthesis of proanthocyanidins, phytate, raffinose family oligosaccharides, vicine, and convicine. Plant Genome 8, (2015).

12. Khazaei, H. et al. Development and validation of a robust, breeder-friendly molecular marker for the *vc- locus in faba bean*. Mol. Breed. 37, 140 (2017).

13. Khazaei, H. et al. Flanking SNP markers for vicine–convicine concentration in faba bean (*Vicia faba* L.). Mol. Breed. 35, 38 (2015).

14. O’Sullivan, D. M. & Angra, D. Advances in faba bean genetics and genomics. Front. Genet. 7, 150 (2016).

15. Kereszt, A. et al. *Agrobacterium rhizogenes*-mediated transformation of soybean to study root biology. Nat. Protoc. 2, 948–952 (2007).

16. Reid, D. et al. Cytokinin biosynthesis promotes cortical cell responses during nodule development. Plant Physiol. 175, 361–375 (2017).

17. Hiltunen, H.-M., Illarionov, B., Hedtke, B., Fischer, M. & Grimm, B. *Arabidopsis* RIBA proteins: two out of three isoforms have lost their bifunctional activity in riboflavin biosynthesis. Int. J. Mol. Sci. 13, 14086–14105 (2012).

18. Brown, E. G. & Roberts, F. M. Formation of vicine and convicine by *Vicia faba*. Phytochemistry 11, 3203–3206 (1972).

19. Frelin, O. et al. A directed-overflow and damage-control N-glycosidase in riboflavin biosynthesis. Biochem. J 466, 137–145 (2015).

20. Lehmann, M. et al. Biosynthesis of riboflavin. Screening for an improved GTP cyclohydrolase II mutant. FEBS J. 276, 4119–4129 (2009).

21. Spoonamore, J. E. & Bandarian, V. Understanding functional divergence in proteins by studying intragenomic homologues. Biochemistry 47, 2592–2600 (2008).

22. Yadav, S. & Karthikeyan, S. Structural and biochemical characterization of GTP cyclohydrolase II from *Helicobacter pylori* reveals its redox dependent catalytic activity. J. Struct. Biol. 192, 100–115 (2015).

## Method References

1. Grabherr, M. G. et al. Full-length transcriptome assembly from RNA-Seq data without a reference genome. Nat. Biotechnol. 29, 644–652 (2011).

2. Fu, L., Niu, B., Zhu, Z., Wu, S. & Li, W. CD-HIT: accelerated for clustering the next-generation sequencing data. Bioinformatics 28, 3150–3152 (2012).

3. Gilbert, D. Gene-omes built from mRNA-seq not genome DNA. Poster presented at the 7th Annual Arthropod Genomics Symposium in Notre Dame, Indiana (2013).

4. Li, H. Aligning sequence reads, clone sequences and assembly contigs with BWA-MEM. 1303.3997 [q-bio.GN] (2013).

5. Waterhouse, R. M. et al. *BUSCO applications from quality assessments to gene prediction and* phylogenomics. Mol. Biol. Evol. 35, 543–548 (2018).

6. Langmead, B. & Salzberg, S. L. Fast gapped-read alignment with Bowtie 2. Nat. Methods 9, 357–359 (2012).

7. Li, B. & Dewey, C. N. RSEM: accurate transcript quantification from RNA-Seq data with or without a reference genome. BMC Bioinformatics 12, 323 (2011).

8. Tautenhahn, R., Patti, G. J., Rinehart, D. & Siuzdak, G. XCMS Online: a web-based platform to process untargeted metabolomic data. Anal. Chem. 84, 5035–5039 (2012).

9. Saeed, A. I. et al. TM4: a free, open-source system for microarray data management and analysis. Biotechniques 34, 374–378 (2003).

10. Li, J., Witten, D. M., Johnstone, I. M. & Tibshirani, R. Normalization, testing, and false discovery rate estimation for RNA-sequencing data. Biostatistics 13, 523–538 (2012).

11. R Core Team. R Foundation for Statistical Computing, Vienna, Austria. R: A language and environment for statistical computing. Available online at https://www.R-project.org/ (2018)

12. Khazaei, H. et al. Flanking SNP markers for vicine–convicine concentration in faba bean (*Vicia faba* L.). Mol. Breed. 35, 38 (2015).

13. Webb, A. et al. A SNP-based consensus genetic map for synteny-based trait targeting in faba bean (*Vicia faba* L.). Plant Biotechnol. J. 14, 177–185 (2016).

14. Broman, K. W., Wu, H., Sen, S. & Churchill, G. A. R/qtl: QTL mapping in experimental crosses. Bioinformatics 19, 889–890 (2003).

15. Weber, E., Engler, C., Gruetzner, R., Werner, S. & Marillonnet, S. A modular cloning system for standardized assembly of multigene constructs. PLoS One 6, e16765 (2011).

16. Nadzieja, M., Stougaard, J. & Reid, D. A Toolkit for high resolution imaging of cell division and phytohormone signaling in legume roots and root nodules. Front. Plant Sci. 10, 1000 (2019).

17. Stougaard, J. *Agrobacterium rhizogenes* as a vector for transforming higher plants. Application in Lotus corniculatus transformation. Methods Mol. Biol. 49, 49–61 (1995).

18. Kereszt, A. et al. *Agrobacterium rhizogenes*-mediated transformation of soybean to study root biology. Nat. Protoc. 2, 948–952 (2007).

19. Armenteros, J. J. A. et al. Detecting sequence signals in targeting peptides using deep learning. Life science alliance 2, (2019).

20. Hiltunen, H.-M., Illarionov, B., Hedtke, B., Fischer, M. & Grimm, B. *Arabidopsis* RIBA proteins: two out of three isoforms have lost their bifunctional activity in riboflavin biosynthesis. Int. J. Mol. Sci. 13, 14086–14105 (2012).

21. Lehmann, M. et al. Biosynthesis of riboflavin. Screening for an improved GTP cyclohydrolase II mutant. FEBS J. 276, 4119–4129 (2009).

