## Supplementary Files 1-7. for "VC1 catalyzes a key step in the biosynthesis of vicine from GTP in faba bean": Supplementary File 5.R

```
### Expression data analysis
{
  ## Load expression data
  setwd("[working directory")
  d <- read.delim("[RNA-seq data]")
  colnames(d)
  head(d)
  
  ## Normalize using PoissonSeq
  library(PoissonSeq)
  norm.facs.pss <- PS.Est.Depth(d[,2:ncol(d)], iter=5, ct.sum=5, ct.mean=0.2)
  norm.facs.pss
  d.norm.pss <- t(t(d[,2:ncol(d)])/norm.facs.pss) 
  d <- cbind(as.data.frame(d[,1]), d.norm.pss)
  colnames(d)[1] <- "Name"
  head(d)
  
  # make function to compute row variance
  RowVar <- function(x) {
    rowSums((x - rowMeans(x))^2)/(dim(x)[2] - 1)
  }
  
  # For each gene, calculate variance across all samples 
  # and across replicates for each sample
  colnames(d)
  d$tot.var <- RowVar(d[,2:27]) # all samples
  d$sam1.var <- RowVar(d[,2:4]) # flower
  d$sam2.var <- RowVar(d[,5:7]) # stem
  d$sam3.var <- RowVar(d[,8:10]) # young leaf
  d$sam4.var <- RowVar(d[,11:13]) # mature leaf
  d$sam5.var <- RowVar(d[,14:16]) # pods expansion
  d$sam6.var <- RowVar(d[,17:19]) # seeds expansion
  d$sam7.var <- RowVar(d[,20:22]) # pods filling
  d$sam8.var <- RowVar(d[,23:25]) # seeds filling
  d$sam9.var <- RowVar(d[,26:27]) # mix
  
  
  
  
  # filter genes for total variance and read counts
  nrow(d)
  hist(log(d$tot.var,10), breaks=50)
  d.filt <- subset(d, tot.var > 0.10)
  nrow(d.filt)
  # filter for total read counts
  hist(log10(rowSums(d.filt[,2:27])))
  d.filt <- subset(d.filt, rowSums(d.filt[,2:19])> 5)
  nrow(d.filt)
  
  # check expression of VC1
  subset(d.filt, Name=="evgLocus_1250620")
  
  
  # For each gene, generate random groups of "replicates" 
  # and calculate variance for comparison with the observed replicate variance
  k=100     #number of iterations
  rep.num =3     #number of replicates
  i=1
  for (i in 1:k) {
    ran.cols <- sample(ncol(d.filt[,2:27]), rep.num) # sample random columns
    ran.cols <- ran.cols + 1 # adjust index
    colnum <- ncol(d.filt) + 1 # specify index of column to be added
    d.filt[,colnum] <- RowVar(d.filt[,ran.cols]) #add random replicate variance
  }
  head(d.filt)
  summary(d.filt)
  
  # calculate mean of observed replicate variances
  d.filt$mean.rep.var <- (d.filt$sam1.var + d.filt$sam2.var + d.filt$sam3.var + d.filt$sam4.var + d.filt$sam5.var + d.filt$sam6.var + d.filt$sam7.var +d.filt$sam8.var )/8
  
  # calculate mean of random replicate variances
  d.filt$mean.ran.var <- rowMeans(d.filt[,38:137])
  
  # Add ratio between observed replicate and random replicate variances
  d.filt$ratio.var <- log(d.filt$mean.rep.var/d.filt$mean.ran.var,2)
  hist(d.filt$ratio.var, breaks=100)
  
  # filter based on the ratios
  d.filt.sorted <- d.filt[order(d.filt$ratio.var),]
  d.filt.sorted <- subset(d.filt.sorted, ratio.var > -Inf)
  head(d.filt.sorted[,c(2:20,127:129)], 20)
  tail(d.filt.sorted[,c(2:20,127:129)], 20)
  hist(d.filt.sorted$ratio.var)
  d.for.pca.1.rna <- subset(d.filt.sorted, ratio.var < -1)
  nrow(d.for.pca.1.rna)
  head(d.for.pca.1.rna)
  # check expression of VC1
  subset(d.for.pca.1.rna, Name=="evgLocus_1250620")
  
  
  ## PCA analysis
  colnames(d.for.pca.1.rna)
  d.for.pca <- d.for.pca.1.rna[,2:25] # not including the mix of all samples
  d.for.pca <- log(1+d.for.pca)
  #rownames(d.for.pca) <- d.sorted$Name
  head(d.for.pca)
  d.pca <- prcomp(d.for.pca, scale=TRUE)
  par(mfrow=c(1,1))
  plot(d.pca$rotation[,1],d.pca$rotation[,2], type="n")
  text(d.pca$rotation[,1],d.pca$rotation[,2], rownames(d.pca$rotation), cex = 0.6)
  #heatmap(as.matrix(d.for.pca), cexCol=0.5)
  
  # Classical MDS
  # N rows (objects) x p columns (variables)
  # each row identified by a unique row name
  d <- dist(t(d.for.pca)) # euclidean distances between the rows
  fit <- cmdscale(d,eig=TRUE, k=2) # k is the number of dim
  fit # view results
  # plot solution
  x <- fit$points[,1]
  y <- fit$points[,2]
  plot(x, y, xlab="Coordinate 1", ylab="Coordinate 2", 
       main="Metric MDS", type="n")
  text(x, y, labels = colnames(d.for.pca), cex=.5)
  
  ## plot using ggplot
  mds <- as.data.frame(cbind(x,y))
  mds$sample <- c("Flower", "Flower", "Flower", "Stem", "Stem", "Stem", "Leaf.young", "Leaf.young", "Leaf.young", "Leaf.mature", "Leaf.mature", "Leaf.mature", "Pods.expansion", "Pods.expansion", "Pods.expansion", "Seeds.expansion", "Seeds.expansion", "Seeds.expansion", "Pods.filling", "Pods.filling", "Pods.filling", "Seeds.filling.embryo", "Seeds.filling.embryo", "Seeds.filling.embryo")
  mds$tissue <- c("Flower", "Flower", "Flower", "Stem", "Stem", "Stem", "Leaf", "Leaf", "Leaf", "Leaf", "Leaf", "Leaf", "Pods", "Pods", "Pods", "Seeds", "Seeds", "Seeds", "Pods", "Pods", "Pods", "Seeds", "Seeds", "Seeds")
  
  
  ## load libraries
  library(ggplot2)
  
  # set graphics parameters
  alpha = 0.8
  font.size = 18
  point.size = 10
  
  ### simple theme with no gridlines
  theme <- theme(panel.grid.major = element_blank(), panel.grid.minor = element_blank(), axis.text=element_text(size=font.size), legend.text=element_text(size=font.size), axis.title=element_text(size=font.size,face="bold"), legend.title=element_text(size=font.size, face="bold"), legend.background = element_rect(fill="transparent"))
  
  
  ## plot mds colored by sample
  filename <- paste(Sys.Date(),"_FabAtlas_MDS.pdf", sep="")
  pdf(filename, 12, 9)
  pv <- ggplot(data=mds,aes(x=x,y=-y))
  pv + geom_point(aes(x=x,y=y,col=sample),na.rm=TRUE,size=point.size,alpha=alpha, shape=16)  +
    labs(x="PCo1",y="PCo2",color="sample")+ theme
  dev.off()
  
}


### Metabolite data analysis

{
# read in the data
dir.met <- "[working directory]"
setwd(dir.met)

d.met.raw <- read.delim("[Metabolite data]", header = TRUE, row.names = 1)
head(d.met.raw)
d.met.int <- d.met.raw[,5:ncol(d.met.raw)]
colnames(d.met.int)


## PCA analysis
# Metabolite data
{
  
  ## Normalize using PoissonSeq
  library(PoissonSeq)
  norm.facs.pss <- PS.Est.Depth(d.met.int[,1:ncol(d.met.int)], iter=5, ct.sum=5, ct.mean=0.2)
  norm.facs.pss
  d.norm.pss <- t(t(d.met.int[,1:ncol(d.met.int)])/norm.facs.pss) 
  str(d.norm.pss)
  d <- cbind(as.data.frame(rownames(d.met.int)), d.norm.pss)
  colnames(d)[1] <- "Name"
  head(d[,c(1:5)])
  head(d.met.int[,c(1:5)])
  #str(d)
  d <- d[,-1]
  d <- log(1+d)
  head(d[,c(1:5)])
  
  # make function to compute row variance
  RowVar <- function(x) {
    rowSums((x - rowMeans(x))^2)/(dim(x)[2] - 1)
  }
  
  # For each gene, calculate variance across all samples 
  # and across replicates for each sample
  colnames(d)
  d$tot.var <- RowVar(d[,1:54]) # all samples
  d$sam1.var <- RowVar(d[,1:8]) # expansion pods
  d$sam2.var <- RowVar(d[,9:16]) # flower
  d$sam3.var <- RowVar(d[,17:24]) # stem
  d$sam4.var <- RowVar(d[,25:32]) # young leaf
  d$sam5.var <- RowVar(d[,33:38]) # mature leaf
  d$sam6.var <- RowVar(d[,39:42]) # filling pods
  d$sam7.var <- RowVar(d[,43:48]) # filling embryos
  d$sam8.var <- RowVar(d[,49:54]) # expansion seeds
  
  colnames(d[,1:8]) # expansion pods
  colnames(d[,9:16]) # flower
  colnames(d[,17:24]) # stem
  colnames(d[,25:32]) # young leaf
  colnames(d[,33:38]) # mature leaf
  colnames(d[,39:42]) # filling pods
  colnames(d[,43:48]) # filling embryos
  colnames(d[,49:54]) # expansion seeds
  
  
  
  # filter genes for total variance and read counts
  nrow(d)
  colnames(d)
  hist(d$tot.var,10, breaks=50)
  d.filt <- subset(d, tot.var > 0.1)
  nrow(d.filt) # 1328
  #check specific features
  head(d)
  d.filt[row.names(d.filt) %in% "84_(+)", ]

  
  # For each gene, generate random groups of "replicates" 
  # and calculate variance for comparison with the observed replicate variance
  k=100     #number of iterations
  rep.num =7     #number of replicates
  i=1
  for (i in 1:k) {
    ran.cols <- sample(ncol(d.filt[,2:54]), rep.num) # sample random columns
    ran.cols <- ran.cols + 1 # adjust index
    colnum <- ncol(d.filt) + 1 # specify index of column to be added
    d.filt[,colnum] <- RowVar(d.filt[,ran.cols]) #add random replicate variance
  }
  head(d.filt)
  summary(d.filt)
  
  # calculate mean of observed replicate variances
  d.filt$mean.rep.var <- (d.filt$sam1.var + d.filt$sam2.var + d.filt$sam3.var + d.filt$sam4.var + d.filt$sam5.var + d.filt$sam6.var + d.filt$sam7.var +d.filt$sam8.var )/8
  
  # calculate mean of random replicate variances
  d.filt$mean.ran.var <- rowMeans(d.filt[,65:164])
  
  # Add ratio between observed replicate and random replicate variances
  d.filt$ratio.var <- log(d.filt$mean.rep.var/d.filt$mean.ran.var,2)
  hist(d.filt$ratio.var, breaks=100)
  
  #check examples
  d.filt[row.names(d.filt) %in% "84_(+)", ]
  
  
  # filter based on the ratios
  d.filt.sorted <- d.filt[order(d.filt$ratio.var),]
  d.filt.sorted <- subset(d.filt.sorted, ratio.var > -Inf)
  head(d.filt.sorted[,c(2:20,164:166)], 20)
  tail(d.filt.sorted[,c(2:20,164:166)], 20)
  hist(d.filt.sorted$ratio.var)
  d.for.pca.1 <- subset(d.filt.sorted, ratio.var < -1)
  nrow(d.for.pca.1) # 1105
  colnames(d.for.pca.1)
  
  ## PCA analysis
  d.for.pca <- d.for.pca.1[,1:54]
  #d.for.pca <- log(1+d.for.pca)
  #rownames(d.for.pca) <- d.sorted$Name
  head(d.for.pca)
  d.pca <- prcomp(d.for.pca, scale=TRUE)
  par(mfrow=c(1,1))
  plot(d.pca$rotation[,1],d.pca$rotation[,2], type="n")
  text(d.pca$rotation[,1],d.pca$rotation[,2], rownames(d.pca$rotation), cex = 0.6)
  #heatmap(as.matrix(d.for.pca), cexCol=0.5)
  
  # Classical MDS
  # N rows (objects) x p columns (variables)
  # each row identified by a unique row name
  d <- dist(t(d.for.pca)) # euclidean distances between the rows
  fit <- cmdscale(d,eig=TRUE, k=2) # k is the number of dim
  fit # view results
  # plot solution
  x <- fit$points[,1]
  y <- fit$points[,2]
  plot(x, y, xlab="Coordinate 1", ylab="Coordinate 2", 
       main="Metric MDS", type="n")
  text(x, y, labels = colnames(d.for.pca), cex=.5)
  
  ## plot using ggplot

  colnames(d.for.pca[,1:8]) # expansion pods
  colnames(d.for.pca[,9:16]) # flower
  colnames(d.for.pca[,17:24]) # stem
  colnames(d.for.pca[,25:32]) # young leaf
  colnames(d.for.pca[,33:38]) # mature leaf
  colnames(d.for.pca[,39:42]) # filling pods
  colnames(d.for.pca[,43:48]) # filling embryos
  colnames(d.for.pca[,49:54]) # expansion seeds


    
  mds <- as.data.frame(cbind(x,y))
  head(mds)
  mds$sample <- c("Pods.expansion", "Pods.expansion", "Pods.expansion", "Pods.expansion", "Pods.expansion", "Pods.expansion", "Pods.expansion", "Pods.expansion", "Flower", "Flower", "Flower", "Flower", "Flower", "Flower", "Flower", "Flower", "Stem", "Stem", "Stem", "Stem", "Stem", "Stem", "Stem", "Stem", "Leaf.young", "Leaf.young", "Leaf.young", "Leaf.young", "Leaf.young", "Leaf.young", "Leaf.young", "Leaf.young", "Leaf.mature", "Leaf.mature", "Leaf.mature", "Leaf.mature", "Leaf.mature", "Leaf.mature", "Pods.filling", "Pods.filling", "Pods.filling", "Pods.filling", "Seeds.filling.embro", "Seeds.filling.embro", "Seeds.filling.embro", "Seeds.filling.embro", "Seeds.filling.embro", "Seeds.filling.embro", "Seeds.expansion", "Seeds.expansion", "Seeds.expansion", "Seeds.expansion", "Seeds.expansion", "Seeds.expansion")
  mds$tissue <- c("Pods", "Pods", "Pods", "Pods", "Pods", "Pods", "Pods", "Pods", "Flower", "Flower", "Flower", "Flower", "Flower", "Flower", "Flower", "Flower", "Stem", "Stem", "Stem", "Stem", "Stem", "Stem", "Stem", "Stem", "Leaf", "Leaf", "Leaf", "Leaf", "Leaf", "Leaf", "Leaf", "Leaf", "Leaf", "Leaf", "Leaf", "Leaf", "Leaf", "Leaf", "Pods", "Pods", "Pods", "Pods", "Seeds", "Seeds", "Seeds", "Seeds", "Seeds", "Seeds", "Seeds", "Seeds", "Seeds", "Seeds", "Seeds", "Seeds")
  nrow(mds)
  mds.reorder <- mds[c(9:16,33:38,25:32,1:8,39:42,49:54,43:48,17:24),]
  nrow(mds.reorder)
  head(mds.reorder)
  
  
  ## load libraries
  library(ggplot2)
  
  # set graphics parameters
  alpha = 0.8
  font.size = 18
  point.size = 10
  
  ### simple theme with no gridlines
  theme <- theme(panel.grid.major = element_blank(), panel.grid.minor = element_blank(), axis.text=element_text(size=font.size), legend.text=element_text(size=font.size), axis.title=element_text(size=font.size,face="bold"), legend.title=element_text(size=font.size, face="bold"), legend.background = element_rect(fill="transparent"))
  
  
  ## plot mds colored by sample
  filename <- paste(Sys.Date(),"_FabAtlas_MET_MDS.pdf", sep="")
  pdf(filename, 12, 9)
  pv <- ggplot(data=mds.reorder,aes(x=x,y=y))
  pv + geom_point(aes(x=x,y=y,col=sample),na.rm=TRUE,size=point.size,alpha=alpha, shape=16)  +
    labs(x="PCo1",y="PCo2",color="sample")+ theme
  dev.off()
  
}
}


### Gene expression to metabolite correlations
{


# get the normalized and filtered expression data ready for analysis
d.rna <- d.for.pca.1.rna[,2:27]
rownames(d.rna) <- d.for.pca.1.rna$Name
head(d.rna)
d.rna.cor.1 <- d.rna[,c(1:24)]

# get the normalized, filtered and clustered metabolite data ready for analysis
d.met.1 <- d.met.raw
head(d.met.1)
colnames(d.met.1)
colnames(d.rna.cor.1)
subset(d.met.1, Cluster=="Group_103")
d.met.filt <- subset(d.met.1, Single=="ok")
nrow(d.met.filt) #843

# average samples to match the RNA-seq samples
fl1 <- (d.met.filt$FL2_GB6_01_6706+d.met.filt$FL2_GB6_02_6707)/2
fl2 <- (d.met.filt$FL3_GB7_01_6708+d.met.filt$FL3_GB7_02_6709)/2
fl3 <- (d.met.filt$FL4GB8_01_6710+d.met.filt$FL4GB8_02_6711)/2

st1 <- (d.met.filt$ST1_GC1_01_6712 + d.met.filt$ST1_GC1_02_6713)/2
st2 <- (d.met.filt$ST2_GC2_01_6714 + d.met.filt$ST2_GC2_02_6715)/2
st3 <- (d.met.filt$ST4_GC4_01_6719 + d.met.filt$ST4_GC4_02_6720)/2

yl1 <- (d.met.filt$YL2_GA7_01_6690 + d.met.filt$YL2_GA7_02_6691)/2
yl2 <- (d.met.filt$YL3_GA8_01_6692 + d.met.filt$YL3_GA8_02_6693)/2
yl3 <- (d.met.filt$YL4GB1_01_6695 + d.met.filt$YL4GB1_02_6696)/2

ml1 <- (d.met.filt$ML1GB2_01_6697 + d.met.filt$ML1GB2_02_6698)/2
#ml2 <- No matching metabolite data
ml3 <- (d.met.filt$ML4_GB4_01_6701 + d.met.filt$ML4_GB4_02_6702)/2

pe1 <- (d.met.filt$EP1_GC5_01_6721 + d.met.filt$EP1_GC5_02_6722)/2
pe2 <- (d.met.filt$EP2_GC6_01_6723 + d.met.filt$EP2_GC6_02_6724)/2
pe3 <- (d.met.filt$EP4_GC8_01_6728 + d.met.filt$EP4_GC8_01_6728)/2

se1 <- (d.met.filt$ex1_GE7_01_6761 + d.met.filt$ex1_GE7_02_6762 + d.met.filt$ex2_GE8_01_6763 + d.met.filt$ex2_GE8_02_6764 + d.met.filt$ex4_BA1_01_6765 + d.met.filt$ex4_BA1_02_6766)/6

fp2 <- (d.met.filt$FP2_GD5_01_6739 + d.met.filt$FP2_GD5_02_6740)/2
fp3 <- (d.met.filt$FP4_GD6_01_6741 + d.met.filt$FP4_GD6_02_6742)/2

fe2 <- (d.met.filt$FE2_GE4_01_6754 + d.met.filt$FE2_GE4_02_6755)/2
fe3 <- (d.met.filt$FE4GE6_01_6758 + d.met.filt$FE4GE6_02_6759)/2

d.met.cor <- as.data.frame(cbind(fl1, fl2, fl3,
                                 st1, st2, st3,
                                 yl1, yl2, yl3,
                                 ml1, ml3,
                                 pe1, pe2, pe3,
                                 se1,
                                 fp2, fp3,
                                 fe2, fe3))

rownames(d.met.cor) <- rownames(d.met.filt)

head(d.met.cor)
summary(d.met.cor)
str(d.met.cor)


# adjust the RNA matrix to match
colnames(d.rna.cor.1)
nrow(d.rna.cor.1) #23910
se1.rna <- (d.rna.cor.1$Seeds_.expansion._1 + d.rna.cor.1$Seeds_.expansion._2 + d.rna.cor.1$Seeds_.expansion._3) /3
d.rna.cor <- cbind(d.rna.cor.1[,c(1:9,10,12,13:15)], se1.rna, d.rna.cor.1[,c(20,21,23,24)])

# check that the two matrices match
colnames(d.rna.cor)
colnames(d.met.cor)


# calculate the correlations
rna.met.cor <- cor(t(d.rna.cor),t(d.met.cor), method="pearson")

# order according to the vicine metabolite cluster (#35)
colnames(rna.met.cor)
colnames(rna.met.cor)[126]
rna.met.cor.vicine <- rna.met.cor[order(-rna.met.cor[,126]),]


# check the vc1 gene
subset(rna.met.cor, rownames(rna.met.cor) %in% "evgLocus_1250620")
# not very strongly correlated with anything

# make a scatterplot
vc1.rna <- as.data.frame(t(subset(d.rna.cor, rownames(d.rna.cor) %in% "evgLocus_1250620")))
vc1.met <- as.data.frame(t(subset(d.met.cor, rownames(d.met.cor) %in% "89_(+)")))

sample <- c("Flower", "Flower", "Flower", 
            "Stem", "Stem", "Stem", 
            "Leaf.young", "Leaf.young", "Leaf.young", 
            "Leaf.mature", "Leaf.mature", 
            "Pods.expansion", "Pods.expansion", "Pods.expansion", 
            "Seeds.expansion",
            "Pods.filling", "Pods.filling", 
            "Seeds.filling.embryo", "Seeds.filling.embryo")
vc1.rna$sample <- sample
vc1.plot <- cbind(vc1.rna, vc1.met)
colnames(vc1.plot) <- c("evgLocus_1250620", "sample", "vicine")

## load libraries
library(ggplot2)

# set graphics parameters
alpha = 0.8
font.size = 18
point.size = 6

### simple theme with no gridlines
theme <- theme(panel.grid.major = element_blank(), panel.grid.minor = element_blank(), axis.text=element_text(size=font.size), legend.text=element_text(size=font.size), axis.title=element_text(size=font.size,face="bold"), legend.title=element_text(size=font.size, face="bold"), legend.background = element_rect(fill="transparent"))


##  scatter plot colored by sample
filename <- paste(Sys.Date(), "vc1_vs_vicine_all_samples_ggplot.pdf", sep="_")
pdf(filename, width=8, height=5)
pv <- ggplot(data=vc1.plot,aes(x=evgLocus_1250620,y=vicine))
pv + geom_point(aes(x=evgLocus_1250620,y=vicine,col=sample),na.rm=TRUE,size=point.size,alpha=alpha, shape=16)  +
  labs(x="evgLocus_1250620",y="Vicine",color="sample")+ theme
dev.off()


## log version
vc1.plot.log <- log(vc1.plot[,c(1,3)])
vc1.plot.log$sample <- vc1.plot$sample
vc1.plot.log <- subset(vc1.plot.log, evgLocus_1250620 > -Inf)

filename <- paste(Sys.Date(), "vc1_vs_vicine_all_samples_ggplot_log.pdf", sep="_")
pdf(filename, width=8, height=5)
pv <- ggplot(data=vc1.plot.log,aes(x=evgLocus_1250620,y=vicine))
pv + geom_point(aes(x=evgLocus_1250620,y=vicine,col=sample),na.rm=TRUE,size=point.size,alpha=alpha, shape=16)  +
  labs(x="evgLocus_1250620",y="Vicine",color="sample")+ xlim(-6,6)+ theme
dev.off()


# without the embro tissue
d.met.cor.ZeroEmb <- d.met.cor[,c(1:17)]
d.rna.cor.ZeroEmb <- d.rna.cor[,c(1:17)]

# calculate the correlations
rna.met.cor.ZeroEmb <- cor(t(d.rna.cor.ZeroEmb),t(d.met.cor.ZeroEmb), method="pearson")

# order according to the vicine metabolite cluster (#126)
rna.met.cor.ZeroEmb.vicine <- rna.met.cor.ZeroEmb[order(-rna.met.cor.ZeroEmb[,126]),]
head(rna.met.cor.ZeroEmb.vicine[,126], 25)
# check the vc1 gene
subset(rna.met.cor.ZeroEmb, rownames(rna.met.cor.ZeroEmb) %in% "evgLocus_1250620")[126]

# make a scatterplot
gene = "evgLocus_1250620"
vc1.rna.ZeroEmb <- as.data.frame(t(subset(d.rna.cor.ZeroEmb, rownames(d.rna.cor.ZeroEmb) %in% gene)))
vc1.met.ZeroEmb <- as.data.frame(t(subset(d.met.cor.ZeroEmb, rownames(d.met.cor.ZeroEmb) %in% "89_(+)")))

sample.ZeroEmb <- c("Flower", "Flower", "Flower", 
            "Stem", "Stem", "Stem", 
            "Leaf.young", "Leaf.young", "Leaf.young", 
            "Leaf.mature", "Leaf.mature", 
            "Pods.expansion", "Pods.expansion", "Pods.expansion", 
            "Seeds.expansion",
            "Pods.filling", "Pods.filling")
vc1.rna.ZeroEmb$sample <- sample.ZeroEmb
vc1.plot.ZeroEmb <- cbind(vc1.rna.ZeroEmb, vc1.met.ZeroEmb)
colnames(vc1.plot.ZeroEmb) <- c("evgLocus_1250620", "sample", "vicine")

## load libraries
library(ggplot2)

# set graphics parameters
alpha = 0.8
font.size = 18
point.size = 5

### simple theme with no gridlines
theme <- theme(panel.grid.major = element_blank(), panel.grid.minor = element_blank(), axis.text=element_text(size=font.size), legend.text=element_text(size=font.size), axis.title=element_text(size=font.size,face="bold"), legend.title=element_text(size=font.size, face="bold"), legend.background = element_rect(fill="transparent"))


##  scatter plot colored by sample
filename <- paste(Sys.Date(), "vc1_vs_vicine_all_samples_ggplot_NoEmb.pdf", sep="_")
pdf(filename, width=8, height=5)
pv <- ggplot(data=vc1.plot.ZeroEmb,aes(x=evgLocus_1250620,y=vicine))
pv + geom_point(aes(x=evgLocus_1250620,y=vicine,col=sample),na.rm=TRUE,size=point.size,alpha=alpha, shape=16)  +
  labs(x="evgLocus_1250620",y="Vicine",color="sample")+ theme
dev.off()


## log version
vc1.plot.ZeroEmb.log <- log(vc1.plot.ZeroEmb[,c(1,3)])
vc1.plot.ZeroEmb.log$sample <- vc1.plot.ZeroEmb$sample
vc1.plot.ZeroEmb.log <- subset(vc1.plot.ZeroEmb.log, evgLocus_1250620 > -Inf)

filename <- paste(Sys.Date(), "vc1_vs_vicine_all_samples_ggplot_log_NoEmb.pdf", sep="_")
pdf(filename, width=8, height=5)
pv <- ggplot(data=vc1.plot.ZeroEmb.log,aes(x=evgLocus_1250620,y=vicine))
pv + geom_point(aes(x=evgLocus_1250620,y=vicine,col=sample),na.rm=TRUE,size=point.size,alpha=alpha, shape=16)  +
  labs(x="evgLocus_1250620",y="Vicine",color="sample")+ xlim(-6,6)+ theme
dev.off()


## average the correlations by metabolic clusters
# transpose the matrix
rna.met.cor.ZeroEmb.t <- t(rna.met.cor.ZeroEmb)
head(rownames(rna.met.cor.ZeroEmb.t))
head(rna.met.cor.ZeroEmb.t[,c(1:5)])

# merge with the original input matrix containing the cluster ids
rna.met.cor.ZeroEmb.t.clusters <- merge(d.met.raw, rna.met.cor.ZeroEmb.t, by="row.names")
head(colnames(rna.met.cor.ZeroEmb.t.clusters),100)
rna.met.cor.ZeroEmb.t.clusters.slim <- rna.met.cor.ZeroEmb.t.clusters[,c(4,60:ncol(rna.met.cor.ZeroEmb.t.clusters))]
head(colnames(rna.met.cor.ZeroEmb.t.clusters.slim),5)
nrow(rna.met.cor.ZeroEmb.t.clusters)
nrow(rna.met.cor.ZeroEmb.t.clusters.slim)
rna.met.cor.ZeroEmb.t.clusters[c(1:5),c(1:8)]
rna.met.cor.ZeroEmb.t.clusters.slim[c(1:5),c(1:8)]
rownames(rna.met.cor.ZeroEmb.t.clusters.slim) <- rna.met.cor.ZeroEmb.t.clusters$Row.names
head(rownames(rna.met.cor.ZeroEmb.t.clusters.slim))
subset(rna.met.cor.ZeroEmb.t.clusters.slim, Cluster=="Group_103")[1:5]


# Aggregate by cluster 
rna.met.cor.ZeroEmb.t.clusters.slim.agg <- aggregate(rna.met.cor.ZeroEmb.t.clusters.slim,
                      by = list(rna.met.cor.ZeroEmb.t.clusters.slim$Cluster),
                      FUN = mean)
head(rna.met.cor.ZeroEmb.t.clusters.slim[,(1:5)])
head(rna.met.cor.ZeroEmb.t.clusters.slim.agg[,(1:5)])
str(rna.met.cor.ZeroEmb.t.clusters.slim.agg)


# transpose the cluster
rna.met.cor.ZeroEmb.t.clusters.slim.agg.t <- as.data.frame(t(rna.met.cor.ZeroEmb.t.clusters.slim.agg),stringsAsFactors = FALSE)
str(rna.met.cor.ZeroEmb.t.clusters.slim.agg.t)
colnames(rna.met.cor.ZeroEmb.t.clusters.slim.agg.t) <- rna.met.cor.ZeroEmb.t.clusters.slim.agg.t[1,]
rna.met.cor.ZeroEmb.t.clusters.slim.agg.t <- rna.met.cor.ZeroEmb.t.clusters.slim.agg.t[-1,]
rna.met.cor.ZeroEmb.t.clusters.slim.agg.t[] <- lapply(rna.met.cor.ZeroEmb.t.clusters.slim.agg.t, type.convert, as.is = TRUE)
str(rna.met.cor.ZeroEmb.t.clusters.slim.agg.t)

head(rna.met.cor.ZeroEmb.t.clusters.slim.agg.t[,(1:5)])
colnames(rna.met.cor.ZeroEmb.t.clusters.slim.agg.t)
head(rownames(rna.met.cor.ZeroEmb.t.clusters.slim.agg.t))
length(rownames(rna.met.cor.ZeroEmb.t.clusters.slim.agg.t))
nrow(rna.met.cor.ZeroEmb.t.clusters.slim.agg.t)

# order according to the vicine metabolite cluster (#2 Group_103)
rna.met.cor.ZeroEmb.t.clusters.slim.agg.t.vicine <- rna.met.cor.ZeroEmb.t.clusters.slim.agg.t[order(-rna.met.cor.ZeroEmb.t.clusters.slim.agg.t[,2]),]
nrow(rna.met.cor.ZeroEmb.t.clusters.slim.agg.t.vicine)
top25.values <- head(rna.met.cor.ZeroEmb.t.clusters.slim.agg.t.vicine$Group_103, 25)
top25.genes <- head(rownames(rna.met.cor.ZeroEmb.t.clusters.slim.agg.t.vicine),25)
top25 <- as.data.frame(cbind(top25.genes, top25.values))
# check the vc1 gene
subset(rna.met.cor.ZeroEmb.t.clusters.slim.agg.t.vicine, rownames(rna.met.cor.ZeroEmb.t.clusters.slim.agg.t.vicine) %in% "evgLocus_1250620")[2]

# make a scatterplot
gene = "evgLocus_1250620"
vc1.rna.ZeroEmb <- as.data.frame(t(subset(d.rna.cor.ZeroEmb, rownames(d.rna.cor.ZeroEmb) %in% gene)))
vc1.met.ZeroEmb <- as.data.frame(t(subset(d.met.cor.ZeroEmb, rownames(d.met.cor.ZeroEmb) %in% "108_(+)")))
plot(t(vc1.rna.ZeroEmb) , t(vc1.met.ZeroEmb))
plot(t(vc1.rna.ZeroEmb) , t(vc1.met.ZeroEmb), log="xy")


## generate a heatmap for the genes showing top correlation with cluster 35
# subset for heatmap
cor.heat <- head(rna.met.cor.ZeroEmb.t.clusters.slim.agg.t.vicine, 20000)
#cor.heat <- rna.met.cor.ZeroEmb.t.clusters.slim.agg.t.vicine
rownames(cor.heat)
# visualise in a heatmap
{
  library(gplots)
  library(viridis)
  
  head(cor.heat)
  
  # filename <- paste(Sys.Date(), "heatmap_cor.pdf", sep="_")
  filename <- paste(Sys.Date(), "heatmap_cor.png", sep="_")
  png(filename, res = 200, width = 4000, height = 4000)
  #pdf(filename, 20 , 20) 
  col.scale <- viridis(14)
  heatmap.2(t(cor.heat), 
            col=col.scale,
            Rowv = TRUE,
            Colv = TRUE,
            trace="none", 
            dendrogram = "none",
            scale= NULL,
            labRow =,
            labCol =
  )
  dev.off()
}


# visualize as heatmaps
vc1.rna <- as.data.frame(t(subset(d.rna.cor, rownames(d.rna.cor) %in% rownames(cor.heat))))
vc1.rna.heatmap <- subset(vc1.rna, !(rownames(vc1.rna) %in% c("Leaf_.young._1",  
                                                              "Leaf_.young._2",  
                                                              "Leaf_.young._3",  
                                                              "Leaf_.mature._1",  
                                                              "Leaf_.mature._3"
                                                              )))
rownames(vc1.rna)
rownames(vc1.rna.heatmap)

filename <- paste(Sys.Date(), "candidate_heatmap.pdf", sep="_")
vc1.rna.heatmap.log <- log(vc1.rna.heatmap)
vc1.rna.heatmap.log[vc1.rna.heatmap.log < -10] <- NA
vc1.rna.heatmap.sqrt <- (vc1.rna.heatmap)^(1/3)
rownames(vc1.rna.heatmap.sqrt)
vc1.rna.heatmap.sqrt.ordered <- vc1.rna.heatmap.sqrt[c(10, 13:14, 11:12, 4:6, 7:9, 1:3),]
pdf(filename, width=10, height=6)
heatmap.2(as.matrix(vc1.rna.heatmap.sqrt.ordered), # add the smallest number larger than zero
          col=col.scale,
          Rowv = FALSE,
          Colv = TRUE,
          trace="none", 
          dendrogram = "column",
          scale= "none",
          labRow =,
          labCol =)
dev.off()


}
```
