## Supplementary Files 1-7. for "VC1 catalyzes a key step in the biosynthesis of vicine from GTP in faba bean": Supplementary File 6.docx

**Suplementary file 6**. Hedin/2 *VC1* and Mélodie /2 *vc1* cDNA sequences and their predicted amino acid sequences. The additional AT dinucleotide in *vc1* and resulting frame shift in the amino acid sequence of vc1 are marked with red font. * indicates premature stop codon.

*VC1* cDNA Hedin/2

ATGGCAGCTGCTACTTTCAATCTCTCTTCTTCCTCCTCAAAGCCACTCTCATCTTCCCGGCTATCCAACCACTTCAAAATTTTCAATGATTTACCTCCTGCGAGACTCAATTATAAATTTGCTTCAACTCATAATGCTGCTTTTAAGGTTAAAGCTGCATTGGTATCTGGAGGGGGTAAAGATCTGGCTGTTGTTGATGAATCCAAGTCGGGGATAACAAACCTTGTTTTCCATGTTGACAACAATGAATTTGATTTGGACAGTCCTGCAGAAGGTTTCGCTTCTATCCCTGAAGCCATTGAAGACATTCGCCAGGGAAAGATGGTAGTAGTTGTAGACGATGAAGACAGAGAAAATGAAGGAGACTTGATAATGGCAGCACAGTTGGCAACACCCGAAGCTATGGCTTTTATAGTGAAGCATGGAAGTGGCATAGTTTGTGTAAGCATGAAAGAGGAAGATCTTGATAGATTGGAACTTCCTTTGATGGTGGACAGTAAAGCTAATGCGCAAAAACATTGTACCGCATTCACTGTGTCAGTGGATGCTAAACATGGTACCACCACAGGGGTGTCAGCTCATGACAGGGCAGCTACTGTCTTGGCACTTGCATCTAGAGGTTCAACTCCGGCTGATTTCAACCGACCAGGCCATATTTTCCCAGTAAAATACACTGAAGGTGGTGTCTTAAAGAGAGCCGGACACACAGAAGCTTCAGTCGATCTTACCGTACTTGCTGGTTTGGATCCGGTGTCAGTTCAGTGTGAGATTGTTGATGAAGATGGTTCCATGGCTAGATTGCCTAAGCTTCGCCAGTTTGCCAAGCGTGAGAATTTGAAAATTGTATCTATTGCTGACTTGATAAGATATAGAAGAAAGAGAGACATATTAATAGAACGCTCTTCTGCTGCAAGATTACCTACTCAGTGGGGGAAATTCACATCATATTGTTATAAGTCTCTCTTAGACGGGATTGAGCATATTGCAATGGTTAAGGGTGACCTTGGAGATGGAGAAGATGTTCTTGTTAGGGTACACTCGGAGTGTCTAACTGGAGACATATTTGGATCTGCCAGATGTGACTGTGGAAATCAGCTTGCACTTTCAATGCAGCAGATTGAGGCTACCGGTAGAGGTGTACTTATATATCTCCGCGGACATGAAGGTAGGGGTATTGGATTGGGCCCCAAGCTCCGTGCATATAACCTACAGGATGATGGACGGGATACCGTAGAAGCCAACGAGGAGTTGGGATTGCCTGTTGACTCTAGGGAGTACGGCATTGGTGCACAGATGCTCAGGGATCTAGGTGTTCAATCTATGAAGTTGATGACTAACAATCCAACTAAATATATTGGTCTCAAAGGTTATGGTTTAACTATTACCGGTAGAATCCCACTCTTAACTCTTATCACTTCTGAGAACCAGAGATACTTGGAGACAAAACGTGTCAAAATGGGCCACGTCTATGGCACTGAGCATAACAATGGTGTTAAAATCACTGTTTGA

VC1 amino acid sequence, Hedin/2

MAAATFNLSSSSTKPLSSSRLSNHFKFFNDLPPARLNYKFASTHNAAFKVKAALVSGGGKDLAVVDESKSGITNLVFHVDNNEFDLDSPAEGFASIPEAIEDIRQGKMVVVVDDEDRENEGDLIMAAQLATPEAMAFIVKHGSGIVCASMKEEDLDRLDLPLMVDSKSNAQKLCTAFTVSVDAKHGTTTGVSANDRAATVLALASRGSTPADFNRPGHIFPVKYTEGGVLKRAGHTEASVDLTVLAGLDPVSVQCEIVDEDGSMARLPKLRQFAKRENLKIVSIADLIRYRRKRDILIERSSAARLPTQWGKFTSYCYKSLLDGIEHIAMVKGDLGDGEDVLVRVHSECLTGDIFGSARCDCGNQLALSMQQIEATGRGVLIYLRGHEGRGIGLGPKLRAYNLQDDGRDTVEANEELGLPVDSREYGIGAQMLRDLGVQSMKLMTNNPTKYIGLKGYGLTITGRIPLLTLITSENQRYLETKRVKMGHVYGTEHNNGVKITV

*vc1* cDNA Melodie/2

ATGGCAGCTGCTACTTTCAATCTCTCTTCTTCATCCACAAAGCCACTCTCATCTTCCCGGCTATCCAACCACTTCAAATTTTTCAATGATTTACCTCCTGCGAGACTCAATTATAAATTTGCTTCAACTCATAATGCTGCTTTTAAGGTTAAAGCTGCATTGGTATCTGGAGGGGGTAAAGATCTGGCTGTTGTTGATGAATCCAAGTCGGGGATAACAAACCTTGTTTTCCATGTTGACAACAATGAATTTGATTTGGACAGTCCTGCAGAAGGTTTCGCTTCTATCCCTGAAGCCATTGAAGACATTCGCCAGGGAAAGATGGTAGTAGTTGTAGACGATGAAGACAGAGAAAATGAAGGAGACTTGATAATGGCAGCACAGTTGGCAACACCCGAAGCTATGGCTTTTATAGTGAAGCATGGAAGTGGCATAGTTTGTGTAAGCATGAAAGAGGAAGATCTTGATAGATTGGAACTTCCTTTGATGGTGGACAGTAAAGCTAATGCGCAAAAACATTGTACCGCATTCACTGTGTCAGTGGATGCTAAACATGGTACCACCACAGGGGTGTCAGCTAATGACAGGGCAGCTACTGTCTTGGCACTTGCATCTAGAGGTTCAACTCCGGCTGATTTCAACCGACCAGGCCATATTTTCCCAGTAAAATACACTGAAGGTGGTGTCTTAAAGAGAGCCGGACACACAGAAGCTTCAGTCGATCTTACCGTACTTGCTGGTTTGGATCCGGTGTCAGTTCAGTGTGAGATTGTTGATGAAGATGGTTCCATGGCTAGATTGCCTAAGCTTCGCCAGTTTGCCAAGCGTGAGAATTTGAAAATTGTATCTATTGCTGACTTGATAAGATATAGAAGAAAGAGAGACATATTAATAGAACGCTCTTCTGCAGCAAGATTACCTACTCAGTGGGGGAAATTCACATCATATTGTTATAAGTCTCTCTTAGACGGGATTGAGCATATTGCAATGGTTAAGGGTGACCTTGGAGATGGAGAAGATGTTCTTGTTAGGGTACACTCGGAGTGTCTAACTGGAGACATATTTGGATCTGCCAGAT**AT**GTGACTGTGGAAATCAGCTTGCACTTTCAATGCAGCAGATTGAGGCTACCGGTAGAGGTGTACTTATATATCTCCGCGGACATGAAGGTAGGGGTATTGGATTGGGCCCCAAGCTCCGTGCATATAACCTACAGGATGATGGACGGGATACCGTAGAAGCCAACGAGGAGTTGGGATTGCCTGTTGACTCTAGGGAGTACGGCATTGGTGCACAGATGCTCAGGGATCTAGGTGTTCAATCTATGAAGTTGATGACTAACAATCCAACTAAATATATTGGTCTCAAAGGTTATGGTTTAACTATTACCGGTAGAATCCCACTCTTAACTCTTATCACTTCAGAGAACCAGAGATACTTGGAGACAAAACGTGCCAAAATGGGCCACGTCTATGGCACTGAGCATAACAATGGTGTTAAAATCACTGTTTGA

*vc1* amino acid sequence Melodie/2

MAAATFNLSSSSTKPLSSSRLSNHFKFFNDLPPARLNYKFASTHNAAFKVKAALVSGGGKDLAVVDESKSGITNLVFHVDNNEFDLDSPAEGFASIPEAIEDIRQGKMVVVVDDEDRENEGDLIMAAQLATPEAMAFIVKHGSGIVCVSMKEEDLDRLELPLMVDSKANAQKHCTAFTVSVDAKHGTTTGVSANDRAATVLALASRGSTPADFNRPGHIFPVKYTEGGVLKRAGHTEASVDLTVLAGLDPVSVQCEIVDEDGSMARLPKLRQFAKRENLKIVSIADLIRYRRKRDILIERSSAARLPTQWGKFTSYCYKSLLDGIEHIAMVKGDLGDGEDVLVRVHSECLTGDIFGSARYVTVEISLHFQCSRLRLPVEVYLYISADMKVGVLDWAPSSVHITYRMMDGIP*KPTRSWDCLLTLGSTALVHRCSGI*VFNL*S**LTIQLNILVSKVMV*LLPVESHS*LLSLQRTRDTWRQNVPKWATSMALSITMVLKSLF
