## Supplementary Files 1-7. for "VC1 catalyzes a key step in the biosynthesis of vicine from GTP in faba bean": Supplementary File 7.docx

**Supplementary File 7. Design of the expression constructs used in the study.**

Synthetic DNA sequence coding for N-terminally tagged VC1 lacking the predicted plastid targeting peptide (cTP). The NdeI and HindIII sites used for cloning are shown in bold, and the sequence coding for the 6X-His tag is underlined. A linker region shown in italics was placed in between the His-tag coding region and the start of the *VC1* sequence.

**CATATG**GCGCATCACCACCACCACCAC*AGCAGCGGTCTGGAAGTGCTGTTTCAAGGTCCG*AAGGCGGCGCTGGTTAGCGGTGGCGGTAAAGATCTGGCGGTGGTTGACGAAAGCAAGAGCGGCATCACCAACCTGGTTTTCCACGTGGACAACAACGAGTTTGACCTGGATAGCCCGGCGGAAGGCTTTGCGAGCATTCCGGAGGCGATCGAAGATATTCGTCAGGGTAAAATGGTGGTTGTGGTTGACGATGAGGATCGTGAGAACGAAGGTGACCTGATTATGGCGGCGCAACTGGCGACCCCGGAAGCGATGGCGTTCATCGTTAAACACGGCAGCGGTATTGTGTGCGTTAGCATGAAGGAAGAGGACCTGGATCGTCTGGAGCTGCCGCTGATGGTGGACAGCAAGGCGAACGCGCAAAAACACTGCACCGCGTTTACCGTGAGCGTTGATGCGAAGCACGGCACCACCACCGGTGTTAGCGCGCATGACCGTGCGGCGACCGTGCTGGCGCTGGCGAGCCGTGGCAGCACCCCGGCGGATTTCAACCGTCCGGGTCACATTTTTCCGGTTAAGTACACCGAGGGCGGTGTGCTGAAACGTGCGGGTCACACCGAAGCGAGCGTTGATCTGACCGTGCTGGCGGGTCTGGACCCGGTGAGCGTTCAGTGCGAGATCGTTGACGAAGATGGCAGCATGGCGCGTCTGCCGAAACTGCGTCAATTCGCGAAGCGTGAGAACCTGAAAATCGTGAGCATTGCGGATCTGATTCGTTATCGTCGTAAGCGTGACATCCTGATTGAACGTAGCAGCGCGGCGCGTCTGCCGACCCAGTGGGGCAAGTTTACCAGCTACTGCTATAAAAGCCTGCTGGACGGCATCGAACACATTGCGATGGTTAAAGGTGACCTGGGCGATGGCGAGGACGTGCTGGTTCGTGTGCACAGCGAATGCCTGACCGGCGATATCTTCGGTAGCGCGCGTTGCGACTGCGGTAACCAGCTGGCGCTGAGCATGCAGCAAATCGAGGCGACCGGCCGTGGTGTGCTGATTTACCTGCGTGGTCACGAAGGCCGTGGTATCGGCCTGGGTCCGAAACTGCGTGCGTATAACCTGCAAGACGATGGCCGTGATACCGTTGAGGCGAACGAGGAACTGGGTCTGCCGGTGGACAGCCGTGAATACGGCATTGGTGCGCAGATGCTGCGTGATCTGGGTGTTCAAAGCATGAAGCTGATGACCAACAACCCGACCAAGTACATCGGCCTGAAAGGCTATGGTCTGACCATCACCGGTCGTATTCCGCTGCTGACCCTGATTACCAGCGAGAACCAACGTTACCTGGAAACCAAGCGTGTTAAAATGGGCCACGTGTATGGTACCGAGCACAACAACGGTGTTAAGATCACCGTGTAA**AAGCTT**

Expression unit of the VC1 complementation vector. i) *Italics* – LjUbi promoter, ii) **bold** – Hedin/2 *VC1* coding sequence with 2 silent mutations removing BpiI/BsaI sites, iii) underline – CaMV 35S terminator.

*GGAGAGAGGATTTTGAGGAAATAATTAATTGAATTACTTGTATTATTGATAAAGTAATTTAGATAAGTTGTTAGTACAACTTATTGCAACTATGGATAGACAAAAATCACTTATTTTAAGGGGAGAAGTATATAACAACTTATTATAAAATTTCTGAGCAGCCGGCCTCCTCCAATCAATCATAGAGAGTGGAGCCCATTCTGGAAAACCAAGGAACCCCCAACTTGCACTGGTGCGGTGGCCCAATTCAAAAAAAACGGGGCCCAGCAAAGTAACGCCGTTGTAACGATTTATCAATCCAAACTCAAAAGGCGGCGAGATGCGTTTAACTCCGTGAAATTAACAAACCGCCAACAACTTGCAATTTGCAACTACCGTTTCCAAGAAGAACTCAACCACACAACGTATCCTATCCCAAACCACACGCAACTAGTGACGCGTCATAAGGACACGTGTCACAATTTGACTGGTTAATAATTTCACCGCTTTTGCTATAAATTACCTCCAATCCCCTTAGCTTCTTCACAATTCAGTTCCCAACCCTAACAATTCTTGTTCATATCGCTTCTCTCTACTTTCAAGGTATGATCCAATTTCTCTCTTCTTTCTCTGTAATCCTTTCGTTGAGTTTTGTTTCCGATCAATCATAGGTAGTTTTCTTGTTTCGAAGCATGAGATCTAGGAATTTTTTGTGATTTTCCAAAATTGAGATCGGTTTGAAATTGAATTTTACAGCTTGAATCTCAGATCTTGTTTTATCAATGTTTTCGATGGCTTGCGATGTAGATCTATGATAATTGTGGTTCAGTTTTGTTAGGAATCGATTTCGGTTTAGCAATTGCAGATTAATTAGGGTTTCCAATTGAATTCTTCAGATCCGTTATGGAATTATGTCAAATAATTTATTCAAATTGGAAATTATTGTTAGATCCACTCTTAATCTGTTTGATCCAAGCTTCAATTAGGGTTTTCACTTGTTTCAATTTCTTGTTATGGATTCTGATTTATCTGTTGATGTTAGATCCACTCTTAATCTGTTTGATCCAAGCGTTAATTAGGGTTTTCACTTGTTTCAATTTCGTGTGTTGGATTCTGATTTATCTGTTGTTGATGTGATTACAGA***ATGATGGCAGCTGCTACTTTCAATCTCTCTTCTTCCTCCTCAAAGCCACTCTCATCTTCCCGGCTATCCAACCACTTCAAAATTTTCAATGATTTACCTCCTGCGAGACTCAATTATAAATTTGCTTCAACTCATAATGCTGCTTTTAAGGTTAAAGCTGCATTGGTATCTGGAGGGGGTAAAGATCTGGCTGTTGTTGATGAATCCAAGTCGGGGATAACAAACCTTGTTTTCCATGTTGACAACAATGAATTTGATTTGGACAGTCCTGCAGAAGGTTTCGCTTCTATCCCTGAAGCCATTGAgGACATTCGCCAGGGAAAGATGGTAGTAGTTGTAGACGATGAgGACAGAGAAAATGAAGGAGACTTGATAATGGCAGCACAGTTGGCAACACCCGAAGCTATGGCTTTTATAGTGAAGCATGGAAGTGGCATAGTTTGTGTAAGCATGAAAGAGGAAGATCTTGATAGATTGGAACTTCCTTTGATGGTGGACAGTAAAGCTAATGCGCAAAAACATTGTACCGCATTCACTGTGTCAGTGGATGCTAAACATGGTACCACCACAGGGGTGTCAGCTCATGACAGGGCAGCTACTGTCTTGGCACTTGCATCTAGAGGTTCAACTCCGGCTGATTTCAACCGACCAGGCCATATTTTCCCAGTAAAATACACTGAAGGTGGTGTCTTAAAGAGAGCCGGACACACAGAAGCTTCAGTCGATCTTACCGTACTTGCTGGTTTGGATCCGGTGTCAGTTCAGTGTGAGATTGTTGATGAAGATGGTTCCATGGCTAGATTGCCTAAGCTTCGCCAGTTTGCCAAGCGTGAGAATTTGAAAATTGTATCTATTGCTGACTTGATAAGATATAGAAGAAAGAGAGACATATTAATAGAACGCTCTTCTGCTGCAAGATTACCTACTCAGTGGGGGAAATTCACATCATATTGTTATAAGTCTCTCTTAGACGGGATTGAGCATATTGCAATGGTTAAGGGTGACCTTGGAGATGGAGAAGATGTTCTTGTTAGGGTACACTCGGAGTGTCTAACTGGAGACATATTTGGATCTGCCAGATGTGACTGTGGAAATCAGCTTGCACTTTCAATGCAGCAGATTGAGGCTACCGGTAGAGGTGTACTTATATATCTCCGCGGACATGAAGGTAGGGGTATTGGATTGGGCCCCAAGCTCCGTGCATATAACCTACAGGATGATGGACGGGATACCGTAGAAGCCAACGAGGAGTTGGGATTGCCTGTTGACTCTAGGGAGTACGGCATTGGTGCACAGATGCTCAGGGATCTAGGTGTTCAATCTATGAAGTTGATGACTAACAATCCAACTAAATATATTGGaCTCAAAGGTTATGGTTTAACTATTACCGGTAGAATCCCACTCTTAACTCTTATCACTTCTGAGAACCAGAGATACTTGGAGACAAAACGTGTCAAAATGGGCCACGTCTATGGCACTGAGCATAACAATGGTGTTAAAATCACTGTTTGA**GCTTCTCTAGCTAGAGTCGATCGACAAGCTCGAGTTTCTCCATAATAATGTGTGAGTAGTTCCCAGATAAGGGAATTAGGGTTCCTATAGGGTTTCGCTCATGTGTTGAGCATATAAGAAACCCTTAGTATGTATTTGTATTTGTAAAATACTTCTATCAATAAAATTTCTAATTCCTAAAACCAAAATCCAGTACTAAAATCCAGAT

Expression unit of the VC1 complementation control vector. i) *Italics* – LjUbi promoter, ii) **bold** – *tYFP-NLS* coding sequence, iii) underline – CaMV 35S terminator.

*GGAGAGAGGATTTTGAGGAAATAATTAATTGAATTACTTGTATTATTGATAAAGTAATTTAGATAAGTTGTTAGTACAACTTATTGCAACTATGGATAGACAAAAATCACTTATTTTAAGGGGAGAAGTATATAACAACTTATTATAAAATTTCTGAGCAGCCGGCCTCCTCCAATCAATCATAGAGAGTGGAGCCCATTCTGGAAAACCAAGGAACCCCCAACTTGCACTGGTGCGGTGGCCCAATTCAAAAAAAACGGGGCCCAGCAAAGTAACGCCGTTGTAACGATTTATCAATCCAAACTCAAAAGGCGGCGAGATGCGTTTAACTCCGTGAAATTAACAAACCGCCAACAACTTGCAATTTGCAACTACCGTTTCCAAGAAGAACTCAACCACACAACGTATCCTATCCCAAACCACACGCAACTAGTGACGCGTCATAAGGACACGTGTCACAATTTGACTGGTTAATAATTTCACCGCTTTTGCTATAAATTACCTCCAATCCCCTTAGCTTCTTCACAATTCAGTTCCCAACCCTAACAATTCTTGTTCATATCGCTTCTCTCTACTTTCAAGGTATGATCCAATTTCTCTCTTCTTTCTCTGTAATCCTTTCGTTGAGTTTTGTTTCCGATCAATCATAGGTAGTTTTCTTGTTTCGAAGCATGAGATCTAGGAATTTTTTGTGATTTTCCAAAATTGAGATCGGTTTGAAATTGAATTTTACAGCTTGAATCTCAGATCTTGTTTTATCAATGTTTTCGATGGCTTGCGATGTAGATCTATGATAATTGTGGTTCAGTTTTGTTAGGAATCGATTTCGGTTTAGCAATTGCAGATTAATTAGGGTTTCCAATTGAATTCTTCAGATCCGTTATGGAATTATGTCAAATAATTTATTCAAATTGGAAATTATTGTTAGATCCACTCTTAATCTGTTTGATCCAAGCTTCAATTAGGGTTTTCACTTGTTTCAATTTCTTGTTATGGATTCTGATTTATCTGTTGATGTTAGATCCACTCTTAATCTGTTTGATCCAAGCGTTAATTAGGGTTTTCACTTGTTTCAATTTCGTGTGTTGGATTCTGATTTATCTGTTGTTGATGTGATTACAGA***ATGGTGAGCAAGGGCGAGGAGCTGTTCACCGGGGTGGTGCCCATCCTGGTCGAGCTGGACGGCGACGTAAACGGCCACAAGTTCAGCGTGTCCGGCGAGGGCGAGGGCGATGCCACCTACGGCAAGCTGACCCTGAAGCTGATCTGCACCACCGGCAAGCTGCCCGTGCCCTGGCCCACCCTCGTGACCACCCTGGGCTACGGCCTGCAGTGCTTCGCCCGCTACCCCGACCACATGAAGCAGCACGACTTCTTCAAGTCCGCCATGCCCGAAGGCTACGTCCAGGAGCGCACCATCTTCTTCAAGGACGACGGCAACTACAAGACCCGCGCCGAGGTGAAGTTCGAGGGCGACACCCTGGTGAACCGCATCGAGCTGAAGGGCATCGACTTCAAGGAGGACGGCAACATCCTGGGGCACAAGCTGGAGTACAACTACAACAGCCACAACGTCTATATCACCGCCGACAAGCAGAAGAACGGCATCAAGGCCAACTTCAAGATCCGCCACAACATCGAGGACGGCGGCGTGCAGCTCGCCGACCACTACCAGCAGAACACCCCCATCGGCGACGGCCCCGTGCTGCTGCCCGACAACCACTACCTGAGCTACCAGTCCGCCCTGAGCAAAGACCCCAACGAGAAGCGCGATCACATGGTCCTGCTGGAGTTCGTGACCGCCGCCGGGATCACTCTCGGCATGGACGAGGCAGCTAGATCCACCATGGTGAGCAAGGGCGAGGAGCTGTTCACCGGGGTGGTGCCCATCCTGGTCGAGCTGGACGGCGACGTAAACGGCCACAAGTTCAGCGTGTCCGGCGAGGGCGAGGGCGATGCCACCTACGGCAAGCTGACCCTGAAGCTGATCTGCACCACCGGCAAGCTGCCCGTGCCCTGGCCCACCCTCGTGACCACCCTGGGCTACGGCCTGCAGTGCTTCGCCCGCTACCCCGACCACATGAAGCAGCACGACTTCTTCAAGTCCGCCATGCCCGAAGGCTACGTCCAGGAGCGCACCATCTTCTTCAAGGACGACGGCAACTACAAGACCCGCGCCGAGGTGAAGTTCGAGGGCGACACCCTGGTGAACCGCATCGAGCTGAAGGGCATCGACTTCAAGGAGGACGGCAACATCCTGGGGCACAAGCTGGAGTACAACTACAACAGCCACAACGTCTATATCACCGCCGACAAGCAGAAGAACGGCATCAAGGCCAACTTCAAGATCCGCCACAACATCGAGGACGGCGGCGTGCAGCTCGCCGACCACTACCAGCAGAACACCCCCATCGGCGACGGCCCCGTGCTGCTGCCCGACAACCACTACCTGAGCTACCAGTCCGCCCTGAGCAAAGACCCCAACGAGAAGCGCGATCACATGGTCCTGCTGGAGTTCGTGACCGCCGCCGGGATCACTCTCGGCATGGACGAGGCAGCTAGATCCACCATGGTGAGCAAGGGCGAGGAGCTGTTCACCGGGGTGGTGCCCATCCTGGTCGAGCTGGACGGCGACGTAAACGGCCACAAGTTCAGCGTGTCCGGCGAGGGCGAGGGCGATGCCACCTACGGCAAGCTGACCCTGAAGCTGATCTGCACCACCGGCAAGCTGCCCGTGCCCTGGCCCACCCTCGTGACCACCCTGGGCTACGGCCTGCAGTGCTTCGCCCGCTACCCCGACCACATGAAGCAGCACGACTTCTTCAAGTCCGCCATGCCCGAAGGCTACGTCCAGGAGCGCACCATCTTCTTCAAGGACGACGGCAACTACAAGACCCGCGCCGAGGTGAAGTTCGAGGGCGACACCCTGGTGAACCGCATCGAGCTGAAGGGCATCGACTTCAAGGAGGACGGCAACATCCTGGGGCACAAGCTGGAGTACAACTACAACAGCCACAACGTCTATATCACCGCCGACAAGCAGAAGAACGGCATCAAGGCCAACTTCAAGATCCGCCACAACATCGAGGACGGCGGCGTGCAGCTCGCCGACCACTACCAGCAGAACACCCCCATCGGCGACGGCCCCGTGCTGCTGCCCGACAACCACTACCTGAGCTACCAGTCCGCCCTGAGCAAAGACCCCAACGAGAAGCGCGATCACATGGTCCTGCTGGAGTTCGTGACCGCCGCCGGGATCACTCTCGGCATGGACGAGCTGTACATTCCTAAGAAGAAGAGAAAGGTTGAGGATTAA**GCTTCTCTAGCTAGAGTCGATCGACAAGCTCGAGTTTCTCCATAATAATGTGTGAGTAGTTCCCAGATAAGGGAATTAGGGTTCCTATAGGGTTTCGCTCATGTGTTGAGCATATAAGAAACCCTTAGTATGTATTTGTATTTGTAAAATACTTCTATCAATAAAATTTCTAATTCCTAAAACCAAAATCCAGTACTAAAATCCAGAT
